# Unsupervised feature computation-based feature selection robustly extracted resting-state functional connectivity patterns related to mental disorders

**DOI:** 10.1101/2025.04.14.648842

**Authors:** Ayumu Yamashita, Takashi Itahashi, Yuki Sakai, Masahiro Takamura, Hiroki Togo, Yujiro Yoshihara, Tomohisa Okada, Hirotaka Yamagata, Kenichiro Harada, Haruto Takagishi, Koichi Hosomi, Naohiro Okada, Osamu Abe, Go Okada, Yasumasa Okamoto, Ryuichiro Hashimoto, Takashi Hanakawa, Toshiya Murai, Koji Matsuo, Hidehiko Takahashi, Kiyoto Kasai, Takuya Hayashi, Shinsuke Koike, Saori C. Tanaka, Mitsuo Kawato, Hiroshi Imamizu, Okito Yamashita, Brain/MINDS Beyond Human Brain MRI Group

**Author notes:** Corresponding author: Ayumu Yamashita.

## Abstract

Research on biomarkers for predicting psychiatric disorders from resting-state functional connectivity (FC) is advancing. While the focus has primarily been on the discriminative performance of biomarkers by machine learning, identification of abnormal FCs in psychiatric disorders has often been treated as a secondary goal. However, it is crucial to investigate the effect size and robustness of the selected FCs because they can be used as potential targets of neurofeedback training or transcranial magnetic stimulation therapy. Here, we incorporated approximately 5,000 runs of resting-state functional magnetic resonance imaging from six datasets, including individuals with three different psychiatric disorders (major depressive disorder [MDD], schizophrenia [SCZ], and autism spectrum disorder [ASD]). We demonstrated that an unsupervised feature-computation-based feature selection method can robustly extract FCs related to psychiatric disorders compared to other conventional supervised feature selection methods. We found that our proposed method robustly extracted FCs with larger effect sizes from the validation dataset compared to different types of feature selection methods based on supervised learning for MDD (Cohen’s d = 0.40 vs. 0.25), SCZ (0.37 vs. 0.28), and ASD (0.17 vs. 0.16). We found 78, 69, and 81 essential FCs for MDD, SCZ, and ASD, respectively, and these FCs were mainly thalamic and motor network FCs. The current study showed that the unsupervised feature-computation-based feature selection method robustly identified abnormal FCs in psychiatric disorders consistently across datasets. The discovery of such robust FCs will contribute to understanding neural mechanisms as abnormal brain signatures in psychiatric disorders. Furthermore, this finding can aid in developing precise therapeutic interventions, such as neurofeedback training or transcranial magnetic stimulation therapy.

## 1. Introduction

Psychiatric disorders constitute a significant global health challenge, affecting millions worldwide (Kessler et al. 2009). Despite extensive research, the neurobiological mechanisms underlying these psychiatric disorders remain largely unknown, thus impeding the development of precise therapeutic interventions (Vogel et al. 2023). The development of machine learning models to discriminate between patients with psychiatric disorders and healthy individuals based on neural bases, such as functional connectivity (FC) measured by resting-state functional magnetic resonance imaging (rs-fMRI), has been remarkable (Woo et al. 2017). Recent national-level and international projects (e.g., Strategic Research Program for Brain Science [SRPBS, https://bicr.atr.jp/decnefpro/], Brain Mapping by Integrated Neurotechnologies for Disease Studies [Brain/MINDS Beyond, https://brainminds-beyond.jp/] (Koike et al. 2021; Tanaka et al. 2021), Autism Brain Imaging Data Exchange [ABIDE, https://fcon_1000.projects.nitrc.org/indi/abide/] (A. Di Martino et al. 2013; Adriana Di Martino et al. 2017), Human Connectome Studies Related To Human Disease [CRHD, https://www.humanconnectome.org/disease-studies] (Glasser, Smith, et al. 2016), the REST-meta-MDD Project [https://rfmri.org/REST-meta-MDD] (Chen et al. 2022) have collected rs-fMRI datasets comprising over a thousand individuals with psychiatric disorders, such as major depressive disorder (MDD), schizophrenia (SCZ), and autism spectrum disorder (ASD). By applying supervised machine learning methods to these large-scale, multi-site datasets, researchers have successfully constructed brain network markers for distinguishing healthy controls (HCs) from individuals with psychiatric disorders based on FC patterns, which can be generalized to independent validation datasets, thereby overcoming the variability in data across imaging sites (Yahata et al. 2016; Ichikawa et al. 2020; Yoshihara et al. 2020; Takagi et al. 2017; A. Yamashita et al. 2020; Kawashima et al. 2024; Drysdale et al. 2017; He, Byrge, and Kennedy 2020; Abraham et al. 2017; Gallo et al. 2023; Itahashi et al. 2024).

Most previous studies have prioritized predicting diagnostic labels, with identifying aberrant neural signatures in psychiatric disorders as a secondary goal. Typically, aberrant FCs are extracted from the weights of predictive machine learning models. However, this approach has two major limitations. First, models optimized for prediction tend to extract FCs with higher signal-to-noise ratios rather than those exhibiting significant differences between HC and psychiatric disorder groups. This makes it difficult to identify FCs that have large effect sizes but high noise levels due to factors, such as imaging site differences. For example, deep brain regions, which are susceptible to noise, may be overlooked. Additionally, diagnostic labels can contain label noise measured as imperfect diagnostic concordance rates (Freedman et al. 2013; Regier et al. 2013), even when based on structured interviews (Shankman et al. 2018; Otsubo et al. 2005; J. B. Williams et al. 1992). Overreliance on these labels may hinder the detection of true neurobiological abnormalities. Second, individual FCs are often unstable (Spisak, Bingel, and Wager 2023; Noble, Scheinost, et al. 2017; Noble, Spann, et al. 2017; Noble, Scheinost, and Constable 2019). Univariate brain-wide association studies (BWAS), which aim to identify single FCs related to individual differences, are particularly problematic due to the high dimensionality and low reliability of single FCs (Marek et al. 2022). To address these issues, we use an unsupervised feature computation-based feature selection method (Taguchi 2017), which extracts single latent spaces represented as linear combinations of multiple FCs, followed by feature selection in these spaces. This approach simultaneously improves reliability and reduces dimensionality.

In this study, we aimed to extract robust and generalizable FCs associated with psychiatric disorders from high-dimensional resting-state FC data using our unsupervised-based feature extraction method. We incorporated approximately 5,000 runs of rs-fMRI from six datasets, including individuals with three different psychiatric disorders (MDD, SCZ, and ASD) and demonstrated that the unsupervised feature computation-based feature selection method can robustly extract individual FCs associated with psychiatric disorders when compared to other conventional supervised feature selection methods. Specifically, we first targeted MDD and assessed the effectiveness of our proposed method by comparing the effect sizes of the extracted FCs with those obtained using conventional supervised-based feature selection methods. We used discovery data and independent validation data from the DecNef SRPB multi-site resting state FC dataset. The discovery data were harmonized using traveling subject data (A. Yamashita et al. 2019), and the validation data were harmonized using ComBat (Fortin et al. 2018; Johnson, Li, and Rabinovic 2007; Fortin et al. 2017; Yu et al. 2018). By using these datasets, we have already succeeded in constructing brain network markers for MDD, SCZ, and ASD that can be generalized across different datasets (A. Yamashita et al. 2020; Kawashima et al. 2024; Itahashi et al. 2024). Next, to investigate the characteristics of the extracted FCs, such as variability across sessions, within a subject, among subjects, imaging sites, and imaging protocols, we utilized another large-scale traveling subject dataset (Koike et al. 2021; O. Yamashita et al. 2024). This dataset is different from the one used for harmonization and was acquired using three imaging protocols from 75 participants who underwent six to eight scans at three or more sites within 6 months. By utilizing the traveling subject dataset, it becomes possible to isolate the various factors described above. Furthermore, to investigate the generalization ability of our proposed method, we applied it to additional datasets, comprising approximately 150 individuals, including 70 with SCZ, from the Center for Biomedical Research Excellence (COBRE), and approximately 2000 individuals, including 1000 with ASD, from the ABIDE dataset.

## 2. Materials and Methods

### 2.1 Ethics statement

All participants included in all datasets provided written informed consent. All recruitment procedures and experimental protocols were approved by the institutional review boards of the principal investigators’ respective institutions (Advanced Telecommunications Research Institute International [approval numbers: 13–133, 14–133, 15–133, 16–133, 17–133, and 18–133], Hiroshima University [E-38], Kyoto Prefectural University of Medicine [RBMR-C-1098], Showa University [B-2014-019 and UMIN000016134], the University of Tokyo [24-496] Faculty of Medicine [3150], Kyoto University [C809 and R0027], and Yamaguchi University [H23-153 and H25-85]).

### 2.2 Participants

We used six rs-fMRI datasets, which were derived from three nationwide projects conducted in Japan (CREST: 2008– 2013, SRPBS: 2008–2018, and BMB: 2018–2024), the COBRE, and ABIDE for the analyses. A comprehensive description of the four datasets from Japan was provided in our previous studies (Koike et al. 2021; Tanaka et al. 2021; A. Yamashita et al. 2020, 2019; O. Yamashita et al. 2024). The participants’ demographics are summarized in Supplementary Tables S1–S4, and data acquisition protocols in each imaging site are described in Supplementary Tables S5–S7. The following is a brief overview of all datasets.

#### (1) The “discovery dataset”

was used to extract the FCs related to psychiatric disorders. This dataset contained rs-fMRI data from 944 participants (564 HCs from 4 sites, 149 individuals with MDD from 3 sites, 106 individuals with SCZ from 3 sites, and 125 individuals with ASD from 2 sites; Supplementary Table S1). Each participant underwent a single open-eye rs-fMRI session that lasted 10 min (Supplementary Table S5). The psychiatric disorders were diagnosed based on DSM-IV at each site. Depression symptoms were evaluated using the Beck Depression Inventory (BDI)-II score obtained from most participants. After excluding any data affected by image processing errors and excessive head motion from further analysis, we included 906 participants (545 HCs, 138 individuals with MDD, 102 individuals with SCZ, and 121 individuals with ASD).

#### (2) The “validation dataset”

was used to validate the effect size of the FCs extracted using the “discovery dataset.” This dataset contained rs-fMRI data from 578 participants (340 HCs, 185 individuals with MDD, and 53 individuals with SCZ from 4 sites; Supplementary Table S2), which are independent samples using different machines and protocols from the discovery dataset. Each participant underwent a single rs-fMRI session lasting 5 or 8 min (Supplementary Table S5). Depressive symptoms were evaluated using the BDI-II score obtained from most participants. After excluding any data affected by image processing errors and excessive head motion from further analysis, we included 571 participants (338 HCs, 181 individuals with MDD, and 52 individuals with SCZ).

#### (3) The “SRPB traveling subject dataset”

was used for the harmonization of the “discovery dataset.” Harmonization is a method for eliminating the effects caused by imaging sites; the details are discussed in our previous research (A. Yamashita et al. 2019). An overview of the harmonization method is provided in the “Control of site differences” section presented below. This dataset contained rs-fMRI data from nine young adult participants (all men; age range: 24–32 years, mean age: 27 ± 2.6 years) who visited 12 sites, participating in fMRI experiments involving two or three runs of 10-min open-eye rs-fMRI within a single experimental session at each site. This dataset was collected using the same imaging protocols as those used for the “discovery dataset.” After excluding any data affected by image processing errors and excessive head motion from further analysis, we included 397 runs from nine participants from 12 imaging sites.

#### (4) The “BMB traveling subject dataset”

was used to investigate the characteristics of the extracted FCs, specifically addressing variability attributed to individual differences (individual factor), intra-participant variability (session factor), scanner factor, and imaging protocol factor, as detailed in our previous research (Koike et al. 2021; O. Yamashita et al. 2024). This dataset contained 1,169 runs of 10-min open-eye rs-fMRI data from 75 participants (48 men and 27 women; mean age: 31.8 ± 10.0 years) from 17 imaging sites. Each participant had visited three or more sites, including one of three hub sites according to a hub-and-spoke model, which is different from the SRPBS traveling subject dataset in which all participants visited all sites. For each participant, data were collected from at least two runs of a 10-min open-eye rs-fMRI task. At least five participants were included at each site.

#### (5) The COBRE

is a publicly available dataset (http://fcon_1000.projects.nitrc.org/indi/retro/cobre.html) that was used as the “SCZ validation dataset” to validate the effect size of the FCs in SCZ. This dataset contained rs-fMRI data from 147 participants (75 HCs and 72 individuals with SCZ; Supplementary Table S3). Each participant underwent a single rs-fMRI session that lasted 6 min (Supplementary Table S5). All participants were screened based on the following exclusion criteria: history of neurological disorder, history of mental retardation, history of severe head trauma with more than 5 min of loss of consciousness, and history of substance abuse or dependence within the last 12 months. Diagnostic information was collected using the Structured Clinical Interview used for DSM Disorders (SCID). After excluding any data affected by image processing errors and excessive head motion from further analysis, we included 75 participants (45 HCs and 30 individuals with SCZ).

#### (6) The ABIDE-I and -II

are publicly available datasets (https://fcon_1000.projects.nitrc.org/indi/abide/) (A. Di Martino et al. 2013; Adriana Di Martino et al. 2017) that were used as “ASD validation datasets” to validate the effect size of FCs in ASD. We integrated and utilized these two datasets. Data from the “ABIDEII-UCLA Long” and “ABIDEII-UPSM Long” were excluded because they were longitudinal in nature. The integrated dataset contained rs-fMRI data from 1941 participants (1091 HCs and 850 individuals with ASD from 33 sites; Supplementary Table S4). Each participant underwent a single rs-fMRI session lasting 5–15 min (Supplementary Table S6 and S7). After excluding any data affected by image processing errors and excessive head motion from further analysis, we included 1643 participants (969 HCs and 674 individuals with ASD).

### 2.3 Preprocessing and calculating the resting state FC matrix

We preprocessed the rs-fMRI data using fmriprep, version 1.0.8 or 1.5.9 (Esteban et al. 2019). The first 10 s of data were discarded to allow for T1 equilibration. Preprocessing steps included slice-timing correction, realignment, coregistration, distortion correction using a field map, segmentation of T1-weighted structural images, and normalization to the Montreal Neurological Institute space. “Fieldmap-less” distortion correction was applied to datasets without field map data. More details on the pipeline are available at http://fmriprep.readthedocs.io/en/latest/workflows.html. Data that were inadequately preprocessed or affected by motions that were too large were excluded from each dataset, respectively.

#### Parcellation of brain regions

To analyze the data using Human Connectome Project (HCP) style surface-based methods, we used ciftify toolbox, version 2.0.2 or 2.3.3 (Dickie et al. 2019). This allowed us to use an HCP-like surface-based pipeline to analyze data that lacked the T2-weighted image required for using HCP pipelines. Next, we used Glasser’s 379 surface-based parcellations (cortical 360 parcellations + subcortical 19 parcellations) as regions of interest (ROIs), considered reliable brain parcellations (Glasser, Coalson, et al. 2016). The blood oxygen level dependent (BOLD) signal time courses were extracted from these 379 ROIs. To facilitate the comparison of our results with those of previous studies, we identified the anatomical names of important ROIs and names of intrinsic brain networks that included the ROIs using anatomical automatic labeling (Tzourio-Mazoyer et al. 2002) and Neurosynth (http://neurosynth.org/locations/).

#### Physiological noise regression

Physiological noise regressors were extracted by applying CompCor (Behzadi et al. 2007). Principal components (PCs) were estimated for the anatomical CompCor (aCompCor). A mask to exclude signals with a cortical origin was obtained by eroding the brain mask and ensuring that it only contained subcortical structures. Five aCompCor components were calculated within the intersection of the subcortical mask and union of the cerebrospinal fluid and white matter masks calculated in the T1-weighted image space after their projection to the native space of functional images in each session. To remove several sources of spurious variance, we used a linear regression with 12 regression parameters, such as six motion parameters, average signals over the whole brain, and five aCompCor components. A temporal bandpass filter was applied to the time series using a first-order Butterworth filter with pass band between 0.01 Hz and 0.08 Hz to restrict the analysis to low-frequency fluctuations, which are characteristic of rs-fMRI BOLD activity (Ciric et al. 2017). Framewise displacement (FD) (Power et al. 2014) was calculated for each functional session using Nipype (https://nipype.readthedocs.io/en/latest/). FD was used in the subsequent scrubbing procedure. To reduce spurious changes in FC resulting from head motion, we removed volumes with FD > 0.5 mm, as proposed in a previous study (Power et al. 2014). FD represents head motion between two consecutive volumes as a scalar quantity (i.e., the summation of absolute displacements in translation and rotation).

FC was calculated as the temporal correlation of rs-fMRI BOLD signals across 379 ROIs in each participant. Several different candidates, such as the tangent method and partial correlation, can be used to measure FC; however, we used Pearson’s correlation coefficient because it is the most commonly used method in previous studies. Fisher’s z-transformed Pearson’s correlation coefficients were calculated between the preprocessed BOLD signal time courses of each possible pair of ROIs and were used to construct 379 × 379 symmetrical connectivity matrices in which each element represents a connection strength between two ROIs. We used 71,631 FC values [(379 × 378)/2] of the lower triangular matrix of the connectivity matrix for further analysis.

#### Control of site differences

Next, we used a traveling subject harmonization method to control for site differences in the FCs in the discovery dataset. This method enabled us to subtract pure site differences (measurement bias) that were estimated from the traveling subject dataset wherein multiple participants travelled to multiple sites (see Supplementary Text S1). We used the ComBat harmonization method (Fortin et al. 2018, 2016; Yu et al. 2018) to control for site differences in the FCs in the independent validation datasets because we did not have a traveling subject data for these datasets. We performed harmonization to correct only for the site difference using information on diagnosis, age, and sex as auxiliary variables in ComBat.

### 2.4 Unsupervised feature computation-based feature selection method

In this section, we explain how important FCs for MDD were extracted. To extract important FCs that distinguish between individuals with MDD and HCs from higher-dimensional FC data (71,631) relative to the number of participants (approximately 1,000), we first performed dimensionality reduction using principal component analysis (PCA). Specifically, within the discovery dataset, we applied PCA to a subset of participants comprising HCs (N = 545) and individuals with MDD (N = 138). Consequently, the FC dimension decreased as many PCs as the number of participants (Feature computation; Fig 1a). PCA scores were calculated for each PC, and we used *t*-tests to detect a PC for which the PCA scores significantly differed between the MDD and HC groups (Fig 1b). Statistical significance was determined using multiple comparison correction, with a false discovery rate (FDR) of *q* set at 0.05 (Benjamini–Hochberg method [FDR-BH method]) (Benjamini and Yekutieli 2001). When a PC with significantly different PCA scores was found, we extracted the FCs with a large contribution to that PC as important FCs for MDD diagnosis (Feature selection; Fig 1c). If we found more than one PC with significantly different PCA scores, we used the PC which had the highest explained variance for simplicity (in our results, this PC also showed the largest *t*-value for the disorder difference). Assuming that the 71,631-dimensional weights follow a normal distribution, we extracted FCs with larger weights as important FCs based on their significantly low occurrence probability after multiple comparisons (FDR-BH method, *q* = 0.05). To investigate the robustness of the chosen FCs, the proposed method was applied to both the “MDD discovery dataset” including 138 individuals with MDD and 545 HCs from the discovery dataset and the “MDD validation dataset” including 185 individuals with MDD and 340 HCs from the validation dataset, and we investigated the extent to which the chosen FCs overlapped.

**Fig 1.**
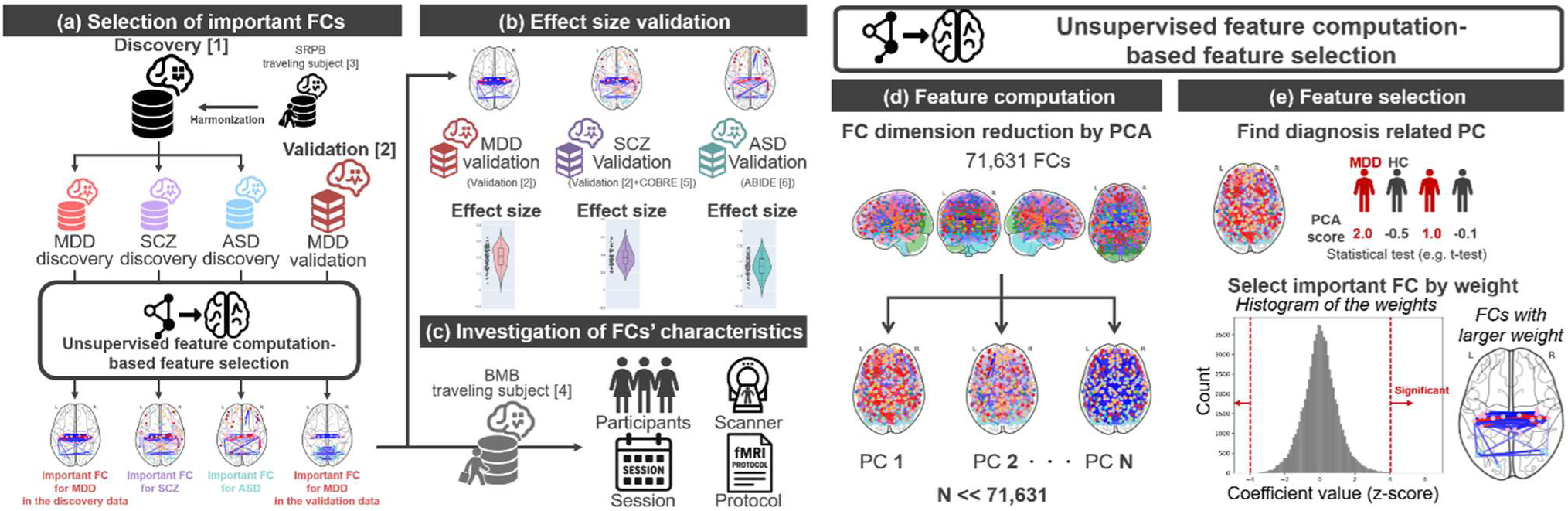
Analysis overview. (a) Important FCs for each disorder are identified using the discovery dataset; for MDD, important FCs are also identified using the validation dataset for investigating robustness of the our method. (b) The effect sizes of the identified FCs are validated using the validation dataset. (c) The characteristics of the identified FCs are investigated using the BMB traveling subject dataset. (d, e) How to extract important FCs for each disorder. (d) Dimension reduction by conducting PCA for all FCs. The color of the nodes represents the network, while the color of the edges indicates the strength of the FC. However, they do not reflect actual data and are for visualization purposes only. (e) Identification of diagnosis related to the PC by calculating the difference in the PC score between HCs and individuals with MDDs. Selection of important FCs by statistical testing of PC weights. FC: functional connectivity, PCA: principal component analysis, HC: healthy control, MDD: major depressive disorder, PC: principal component

To investigate whether the selected PC was MDD-diagnosis specific, we conducted the same feature selection procedure for other factors, such as, age, sex, head motion, and imaging sites. Furthermore, we conducted the same feature selection procedure for depressive symptoms (BDI score). Significant PCs were identified using a two-sample *t*-test for the binary variable, sex. PCs with significant Pearson’s correlations along with PC scores were identified for the continuous variables, the BDI score, FD value, and age. PCs significantly influenced by the imaging sites, a categorical variable, were identified using one-way analysis of variance. The criterion for significance was set at *q* < 0.05 based on the FDR obtained with the FDR-BH method for multiple comparisons correction.

### 2.5 Comparison of robustness in effect sizes of selected FCs with that of other feature-selection methods

Next, we investigated the robustness in effect size (Hedge’s g between the MDD and HC groups) of the selected FCs using the discovery and validation datasets. We considered that if the proposed method robustly selected the FCs with large effect sizes in the discovery dataset, these FCs would also have large effect sizes in the validation dataset. To compare different feature-selection methods, we employed regularization-based feature selection using machine learning methods, such as Least Absolute Shrinkage and Selection Operator (LASSO), and univariate feature selection using a *t*-test with two multiple comparison corrections (FDR-BH and Bonferroni methods). For more detailed information about the regularization-based feature selection method, see Supplementary Text S2 and our previous study (A. Yamashita et al. 2020).

### 2.6 Constructing a network marker using selected FCs

We investigated whether using the FCs selected by our proposed method would improve the prediction performance in MDD diagnosis. We constructed a prediction model using the exactly same method (subsampling and ensemble LASSO) as in our previous research (A. Yamashita et al. 2020). Unsupervised-based feature computation and feature extraction were performed for each subsampling to avoid information leaks, and the performance of the constructed predictive model was tested using a validation dataset (see Supplementary Text S2 for more details).

### 2.7 Validation in other psychiatric disorders

Finally, we tested whether our proposed method could extract important FCs with larger effect sizes without overfitting even in the SCZ and ASD datasets. We applied our proposed method to the “SCZ discovery dataset” including 102 individuals with SCZ and 545 HCs from the discovery dataset. We investigated the effect size of the selected FCs by using “the SCZ validation dataset” including 52 individuals with SCZ and 338 HCs from the validation dataset in addition to 30 individuals with SCZ and 45 HCs from the COBRE dataset. For ASD, we applied our method to a “ASD discovery dataset” including 121 individuals with ASD and 545 HCs from the discovery dataset. We investigated the effect size of the selected FCs by using the ASD validation dataset including 674 individuals with ASD and 969 HCs from the ABIDE-I and ABIDE-II datasets.

## 3. Results

### 3.1 PCA of FCs

We first extracted diagnosis-related PCs by PCA of FCs in the MDD discovery dataset. We investigated the association of PC scores with MDD diagnosis, depressive symptoms (BDI score), age, sex, head motion (FD value), and imaging site. We found that the second PC was significantly associated with MDD diagnosis, seven PCs were significantly associated with age, five PCs were significantly associated with sex, and 12 PCs were significantly associated with imaging sites (Fig 2, visualization of only the top 20 PCs). The PC significantly associated with MDD diagnosis (PC2) was not significantly associated with the other factors. Although no PCs were significantly associated with the BDI score, the PC significantly associated with MDD diagnosis exhibited the strongest correlation with the BDI score. This result indicates that we successfully extracted latent factors solely associated with MDD diagnosis, independent of other factors, such as age, sex, and imaging sites, by using an unsupervised method.

**Fig 2.**
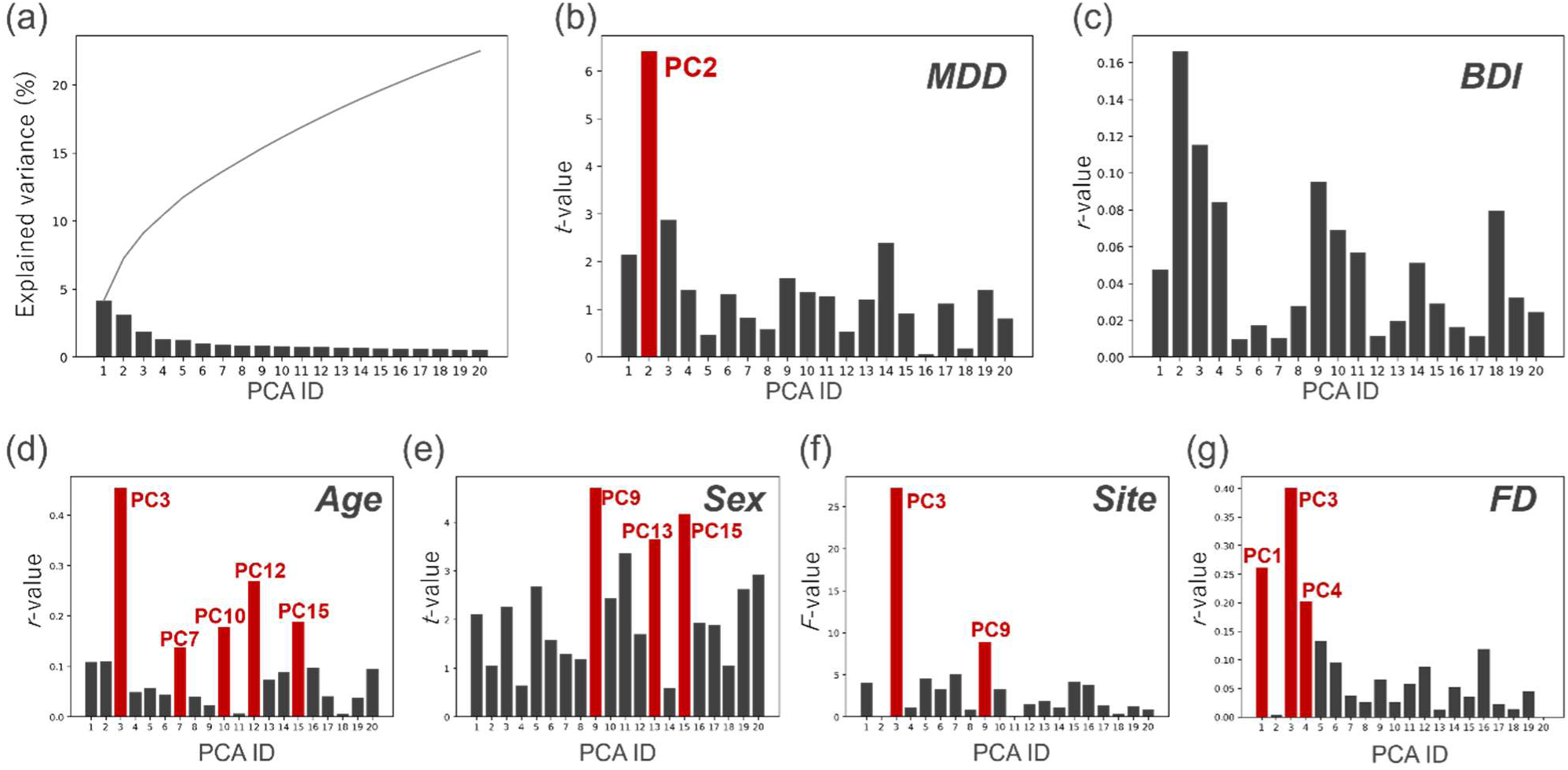
Relationship between the principal component (PC) score and factors in the top 20 PCs. (a) Explained variance of the top 20 PCs and their cumulative summation. (b–g) Relationship between each factor and the top 20 PCs. (b) Difference in PC scores between HCs and individuals with MDDs (*t*-value). (c) Pearson’s correlation coefficients between PC score and BDI score. (d) Pearson’s correlation coefficients between the PC score and age. (e) Difference in PC scores between men and women (*t*-value). (f) Difference in PC scores across imaging sites (*F*-value). (g) Pearson’s correlation coefficients between the PC score and head motion (framewise displacement value). The red bar shows a significant relationship with the factor. Here, we only visualized the top 20 PCs for visualization purpose. Of note, we used all PCs for the analyses. PC: principal component, BDI: Beck Depression Inventory, FD: framewise displacement, PCA: principal component analysis, HC: healthy control, MDD: major depressive disorder

### 3.2 Selected FCs and overlap

We investigated which FCs are crucial for MDD diagnosis by examining the weights of the selected PC. Specifically, we standardized the weights of 71,631 FCs and calculated the probability of occurrence for each weight by assuming that the distribution of weights followed a standard normal distribution. FCs that met the criterion of *q* < 0.05 after multiple comparison correction were identified as significant FCs involved in MDD diagnosis. As a result, we found 78 important FCs (Fig 3 left). The FCs that were primarily involved were bilateral thalamic FCs, thalamo-motor FCs, and interhemispheric motor FCs (see Supplementary Table S8 for more details). Furthermore, we explored whether the same FCs would be selected on applying our proposed method in a validation dataset. We found that the second PC was also significantly associated with MDD diagnosis, and we identified 65 important FCs (Fig 3 right, Supplementary Table S9), of which 20 FCs overlapped with those found in the discovery dataset. Notably, unlike the results in the discovery dataset, the PC also showed significant associations with age in the validation dataset.

**Fig 3.**
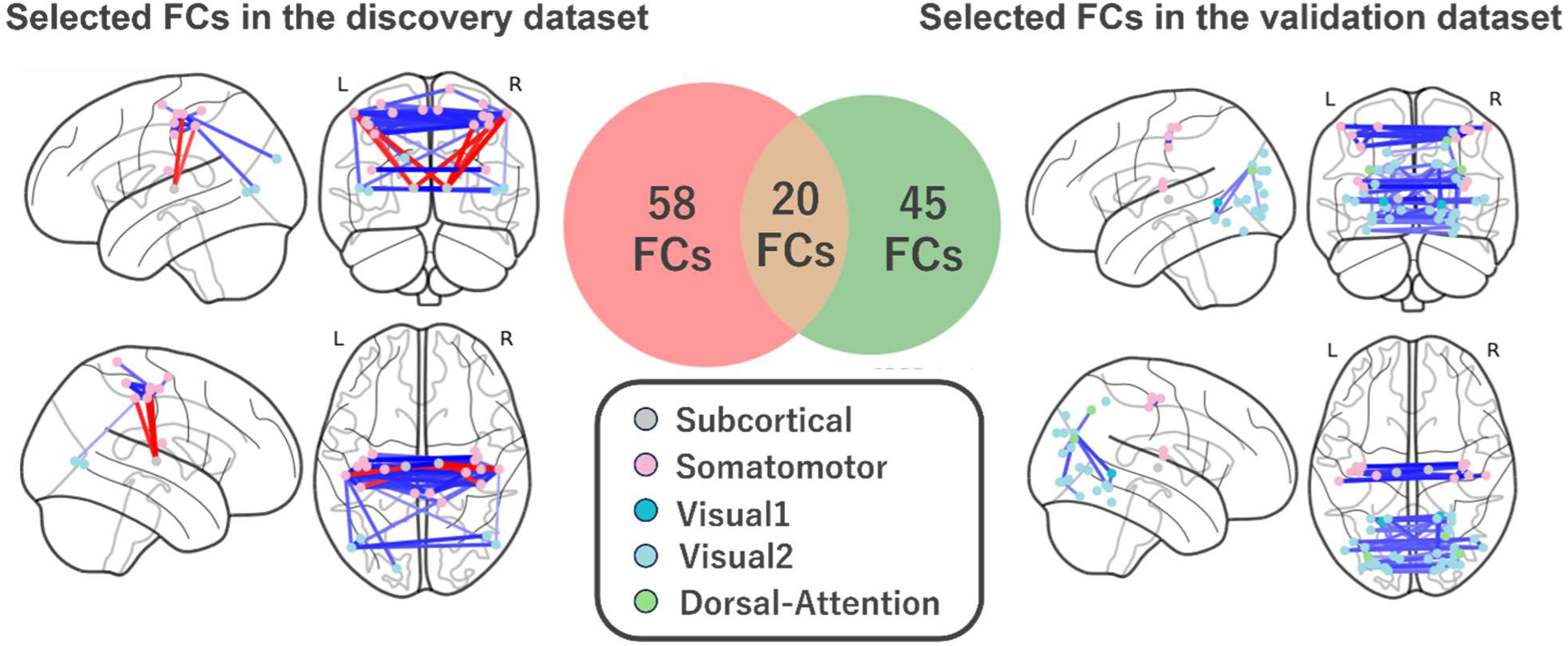
Selected and overlapping FCs. Selected FCs in the discovery dataset (left) and those in the validation dataset (right). Seventy-eight FCs were selected in the discovery dataset, and 65 FCs were selected in the validation dataset. Twenty FCs overlapped between them. Regions are color-coded according to the intrinsic network. The FC state exhibiting a smaller (i.e., more negative) and greater (more positive) mean correlation index in the MDD population than in the HC population is termed under-(blue line) and over-connectivity (red line), respectively. FC: functional connectivity, HC: healthy control, MDD: major depressive disorder

By contrast, on using LASSO, 25 FCs were extracted from the discovery dataset (A. Yamashita et al. 2020), and 65 FCs, from the validation dataset, with no overlapping FCs. That is, the unsupervised-based method exhibited high stability as a FC selection method regardless of which datasets were used, while the supervised learning-based FC selection method exhibited low stability.

### 3.3 Selected FCs had larger effect size with small overfitting

Next, we investigated the effect size (Hedge’s *g* between the MDD and HC groups) of the selected FCs in the discovery and validation datasets. The results showed that the FCs selected by our proposed method had the largest effect size in the validation dataset (Fig 4 and Table 1), while the FCs selected by LASSO had the largest effect size in the discovery dataset. In other words, the unsupervised-based method selected the FCs with a larger effect size regardless of the dataset, while the supervised-based method selected the FCs that were overfitting to the discovery dataset. To visualize the extent of the overfitting of the selected FCs, we plotted Hedge’s *g* from the discovery dataset on the x-axis against Hedge’s *g* from the validation dataset on the y-axis (Fig 5). Specifically, the yellow area in Figure 5 represents FCs with a larger effect size in the discovery dataset than in the validation dataset, suggesting that these FCs are overfitting.

**Fig 4.**
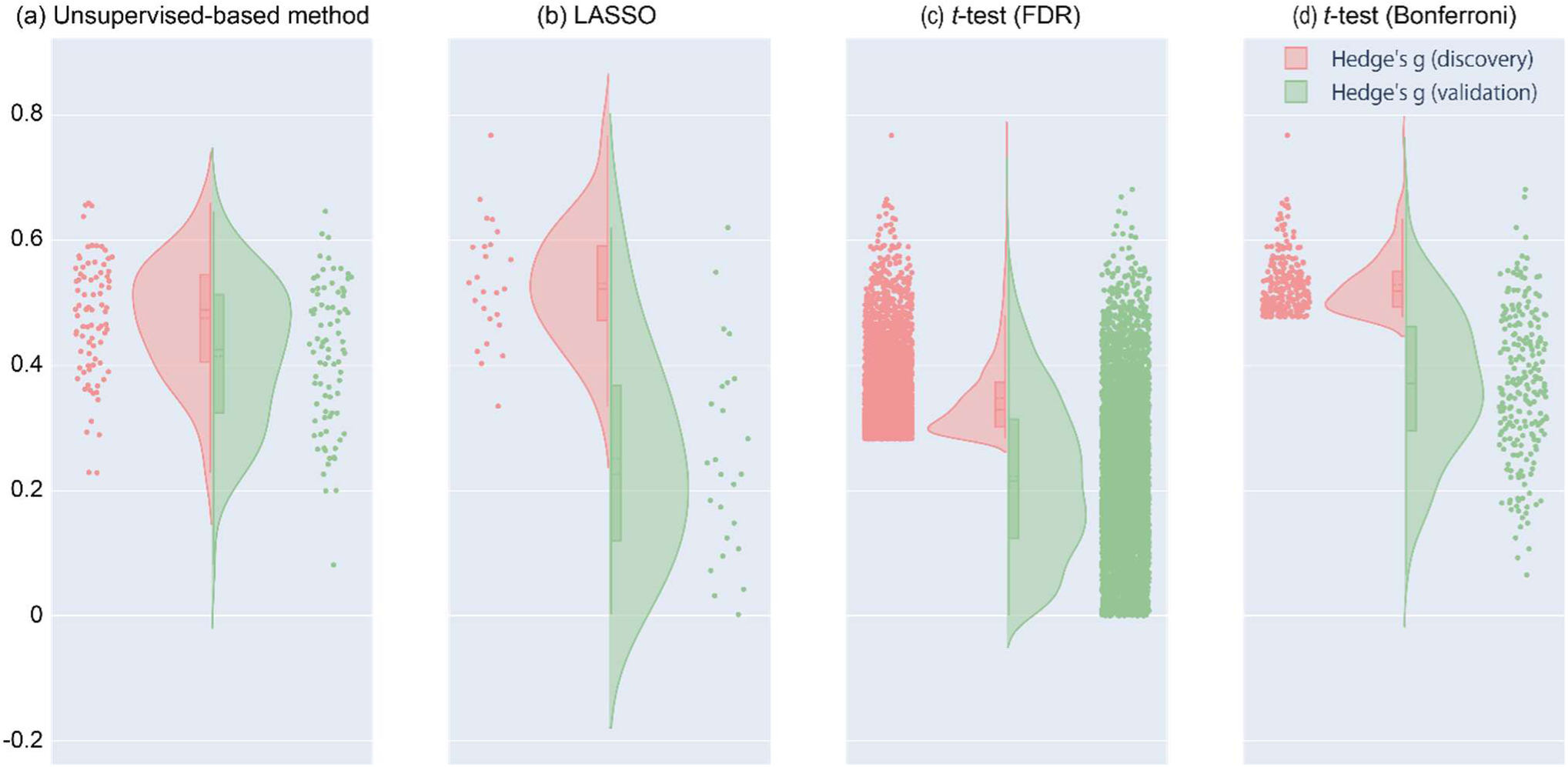
Effect size of selected FCs in both the discovery and validation datasets. (a) FCs selected by the unsupervised-based method. (b) FCs selected by LASSO. (c) FCs selected by a *t*-test using the FDR-BH method. (d) FCs selected by a *t*-test using the Bonferroni method. Each point represents the effect size (Hedge’s *g*) of each FC. The pink points indicate the effect size in the discovery dataset, and the green points indicate the effect size in the validation dataset. Additionally, the histograms for each dataset are also shown. FC: functional connectivity, LASSO: Least Absolute Shrinkage and Selection Operator, FDR: false discovery rate, BH: Benjamini–Hochberg

**Fig 5.**
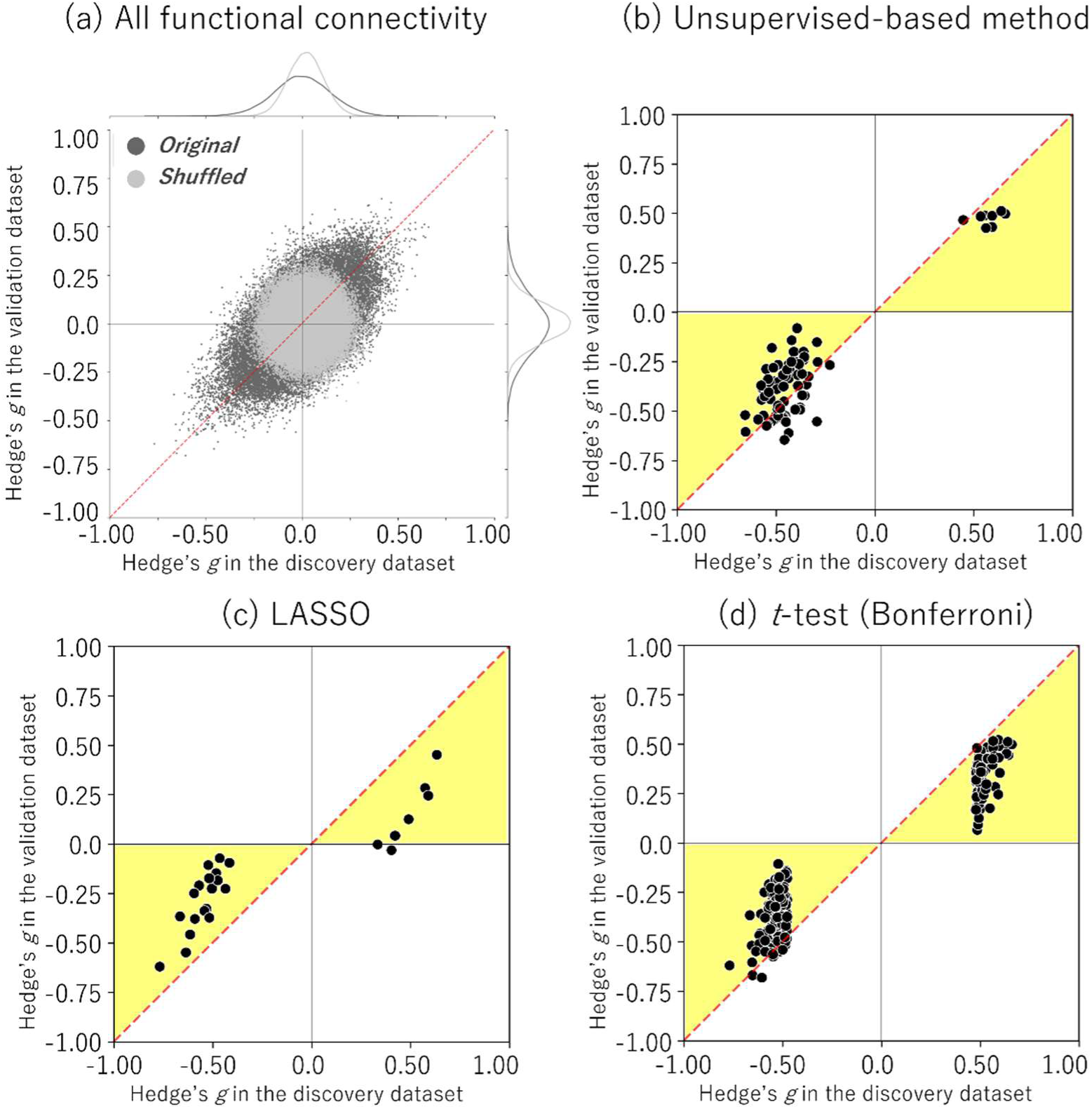
Mass univariate analysis of selected FCs. Reproducibility across the two datasets (discovery and validation) in terms of diagnosis effects (Hedge’s *g*). Scatter plot and histograms of the diagnosis effect size are shown. Each point in the scatter plot represents the diagnosis effect in the discovery dataset in the abscissa and that for the validation dataset in the ordinate for each FC. The original data are shown in dark grey (black), while the shuffled data in which subject information was permuted are shown in gray. (a) All FCs (71,631). (b) FCs selected by the unsupervised-based method. (c) FCs selected by LASSO. (d) FCs selected by a *t*-test using the Bonferroni method. The yellow area represents FCs with a large effect size in the discovery dataset, suggesting that these FCs are overfitting. FC: functional connectivity, LASSO: Least Absolute Shrinkage and Selection Operator

**Table 1.**
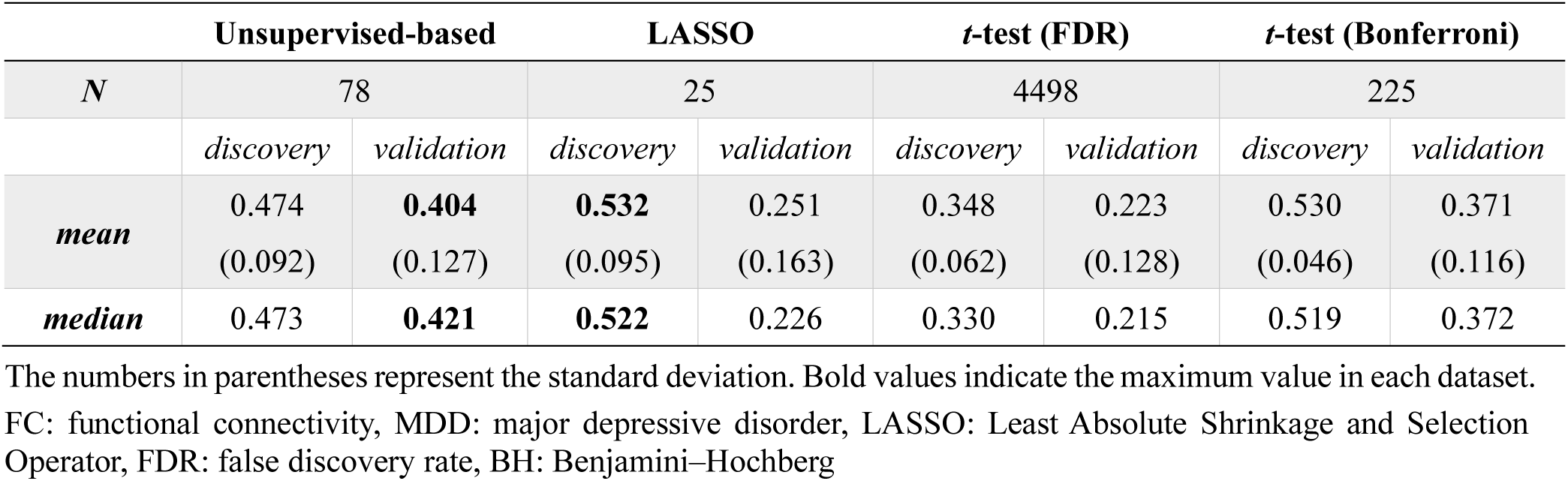
Statistics for selected FCs across methods in MDD.

### 3.4 Network marker construction using only selected FCs

Next, we tested whether using these FCs would improve the prediction performance in MDD diagnosis. The results showed that feature selection using our proposed method did not improve the prediction performance (Table 2). We further tested whether lowering the threshold during feature selection would improve prediction performance. We observed a slight improvement in prediction performance (Area under the curve increased from 0.760 to 0.767, and the Matthews correlation coefficient increased from 0.347 to 0.387) (Table 2).

**Table 2.** Prediction performances of each feature selection method.

|  | AUC | Accuracy | Sensitivity | Specificity | MCC | Average #FC<br>for training |
| --- | --- | --- | --- | --- | --- | --- |
| **Unsupervised-based** | 0.752 | 0.668 | 0.751 | 0.623 | 0.357 | 80 |
| ***t*-test (FDR)** | 0.746 | 0.660 | 0.707 | 0.635 | 0.327 | 615.5 |
| ***t*-test (Bonferroni)** | 0.773 | 0.678 | 0.81 | 0.608 | 0.397 | 19 |
| **No feature selection** | 0.760 | 0.666 | 0.735 | 0.629 | 0.347 | 71631 |
| **Unsupervised-based**<br>**(aggressive)** | 0.768 | 0.680 | 0.779 | 0.626 | 0.387 | 1394.5 |
| ***t*-test (without**<br>**multiple correction)** | 0.761 | 0.674 | 0.740 | 0.638 | 0.361 | 9935.5 |

FC: functional connectivity, FDR: false discovery rate, AUC: area under the curve, MCC: Matthews Correlation Coefficient

Subsequently, we explored why there was no improvement in prediction performance, despite a significant increase in the effect size of the differences between the psychiatric disorder and HC groups in the FCs selected by our proposed method. Our previous research (O. Yamashita et al. 2024) revealed that the reason why we achieved higher prediction performance using subsampling and ensemble LASSO was that the LASSO automatically selected the FCs that had not only a larger effect size between the MDD and HC groups but also a smaller difference among confounding factors, such as site difference and protocol difference. Using the BMB traveling subject dataset (O. Yamashita et al. 2024; Koike et al. 2021), it was possible to divide the factors affecting the variability of FCs into individual differences (individual factor), intra-participant variability (session factor), scanner factor, and imaging protocol factor. Therefore, we investigated how these confounding factors were represented in the FCs selected by each method (see Supplementary Text S3 for more details). The results showed that the FCs selected by our proposed method were significantly larger for all factors than those selected by other methods (Wilcoxon rank-sum test, *p* < 0.01) (Fig 6 and Table 3). This result suggests that the selected FCs identified by the unsupervised-based method inherently reflect multiple factors. In other words, while these FCs exhibit a larger effect size in distinguishing between the psychiatric disorder and HC groups, their direct contribution to prediction may be influenced by these multiple factors. Additionally, it implies that these FCs are unlikely to be chosen when using methods optimized for prediction in supervised machine learning because the supervised method prioritizes prediction performance rather than these effect sizes.

**Fig 6.**
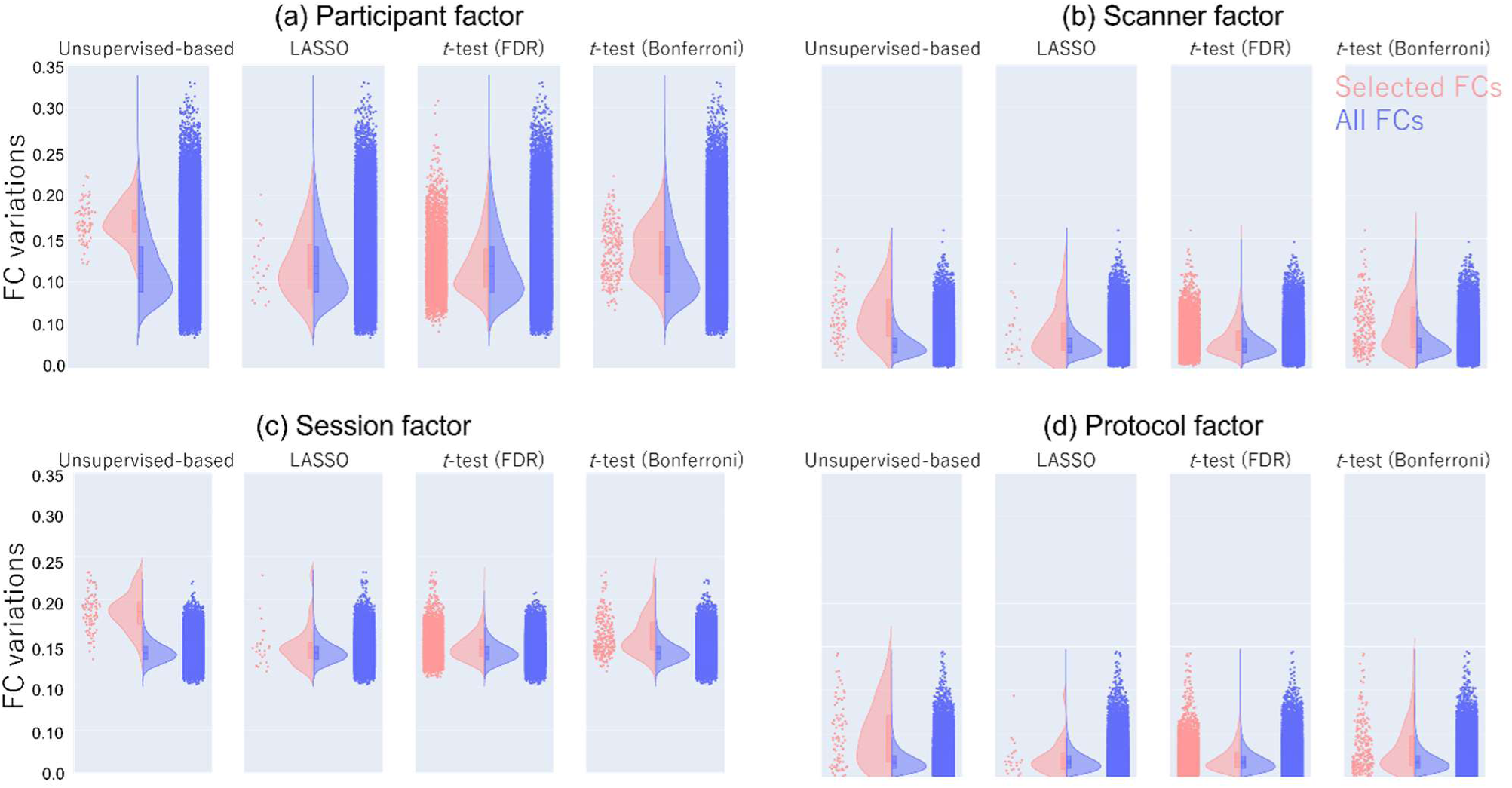
Characteristics of selected FCs. Each point represents the FC variations of each FC. The red points indicate the FC variations in the selected FCs, and the blue points indicate the FC variations in the all FCs. Additionally, the histograms for each dataset are also shown. (a) Participant factor (i.e., individual differences in FC value across traveling subjects). (b) Scanner factor (e.g., difference in FC value between Siemens and GE). (c) Session factor (i.e., differences in FC value across runs within same scanner and same participant. (d) Protocol factor (e.g., difference in FC value between SRPB and HARP protocols). FC: functional connectivity, LASSO: Least Absolute Shrinkage and Selection Operator, FDR: false discovery rate

**Table 3.** Effect of factors on selected FCs in MDD.

|  | Participants | Scanner | Session | Protocol |
| --- | --- | --- | --- | --- |
| **Unsupervised-based** | 0.169 (0.022) | 0.065 (0.030) | 0.186 (0.020) | 0.049 (0.037) |
| **LASSO** | 0.118 (0.034) | 0.042 (0.029) | 0.147 (0.024) | 0.022 (0.019) |
| **t-test (BH-FDR)** | 0.118 (0.033) | 0.034 (0.019) | 0.146 (0.016) | 0.022 (0.016) |
| **t-test (Bonferroni)** | 0.134 (0.032) | 0.050 (0.030) | 0.159 (0.022) | 0.034 (0.029) |
| **All FCs** | 0.118 (0.039) | 0.029 (0.014) | 0.139 (0.012) | 0.019 (0.012) |

The effect was calculated as FC variations within factor (please refer Yamashita O et al., 2024 for more details). The numbers in parentheses represent the standard deviation.
FC: functional connectivity, MDD: major depressive disorder, LASSO: Least Absolute Shrinkage and Selection Operator, BH-FDR: Benjamini–Hochberg method false discovery rate

**Table 4.** Effect size of selected FCs across methods in SCZ.

|  | Unsupervised-based |  | LASSO |  | *t*-test (FDR) |  | *t*-test (Bonferroni) |  |
| --- | --- | --- | --- | --- | --- | --- | --- | --- |
| *N* | 69 |  | 41 |  | 9863 |  | 777 |  |
|  | *discovery* | *validation* | *discovery* | *validation* | *discovery* | *validation* | *discovery* | *validation* |
| ***mean*** | 0.603<br>(0.138) | **0.373**<br>(0.098) | **0.685**<br>(0.168) | 0.261<br>(0.162) | 0.393<br>(0.092) | 0.169<br>(0.11) | 0.619<br>(0.076) | 0.276<br>(0.125) |
| ***median*** | 0.608 | **0.369** | **0.645** | 0.288 | 0.367 | 0.157 | 0.599 | 0.281 |

The numbers in parentheses represent the standard deviation. Bold values indicate the maximum value in each dataset. FC: functional connectivity, SCZ: schizophrenia, LASSO: Least Absolute Shrinkage and Selection Operator, FDR: false discovery rate

**Table 5.** Effect size of selected FCs across methods in ASD.

|  | Unsupervised-based |  | LASSO |  | *t*-test (FDR) |  | *t*-test (Bonferroni) |  |
| --- | --- | --- | --- | --- | --- | --- | --- | --- |
| *N* | 78 |  | 20 |  | 1659 |  | 21 |  |
|  | *discovery* | *validation* | *discovery* | *validation* | *discovery* | *validation* | *discovery* | *validation* |
| ***mean*** | 0.226<br>(0.094) | **0.165**<br>(0.066) | 0.481<br>(0.061) | 0.15<br>(0.081) | 0.374<br>(0.041) | 0.11<br>(0.075) | **0.531**<br>(0.026) | 0.158<br>(0.076) |
| ***median*** | 0.236 | 0.163 | 0.479 | 0.17 | 0.363 | 0.099 | **0.523** | **0.175** |

The numbers in parentheses represent the standard deviation. Bold values indicate the maximum value in each dataset. FC: functional connectivity, ASD, autism spectrum disorder, LASSO: Least Absolute Shrinkage and Selection Operator, FDR: false discovery rate

### 3.5 Validation in other psychiatric disorders

We tested whether our proposed method could extract FCs with larger effect sizes even in the SCZ and ASD discovery datasets. Consequently, we found that two PCs were significantly associated with SCZ diagnosis, and one PC was significantly associated with ASD diagnosis (Supplementary Figures S1b and S2b). These diagnosis-related PCs were not significantly associated with the other factors (age, sex, site and head motion), except for the site factor in the ASD dataset.

Consequently, we found 69 FCs for SCZ and 81 FCs for ASD. Interestingly, these FCs, similar to the MDD-related FCs, were primarily involved in bilateral thalamic FCs, thalamo-motor FCs, and interhemispheric motor FCs (see Supplementary Tables S10 and S11 for more details). The average effect sizes of these FCs in the validation datasets were the highest with our proposed method in both disorders. Statistical analysis showed that the average effect size of SCZ was significantly larger in our proposed method than in the other methods (*t*-test; vs. LASSO: *t* = 4.51, *p* = 1.6 x 10^-5^, dof = 108; vs. *t*-test [BH-FDR]: *t* = 15.36, *p* = 1.23 x 10^-52^, dof = 9930; vs. *t*-test [Bonferroni]: *t* = 6.26, *p* = 6.1 x10^-10^, dof = 844; Figure 7a). By contrast, the average effect size of ASD was significantly larger in our proposed method than in the *t*-test with FDR (*t*-test; vs *t*-test [FDR]: *t* = 6.34, *p* = 2.4 x 10^-10^, dof = 1735); however, it was not significantly different from that in the other methods (*t*-test; vs. LASSO: *t* = 0.83, *p* = 0.41, dof = 96; vs. *t*-test [Bonferroni]: *t* = 0.44, *p* = 0.66, dof = 97; Figure 7b). To investigate the degree to which the effect sizes were similar between the discovery and validation datasets, we calculated Pearson’s correlation coefficients between the *g*-values in the discovery and validation datasets for each disorder and found that the correlations were highest for MDD, followed by SCZ and ASD (*r* = 0.57, 0.52, 0.41, respectively; Figure 7c-e). This result indicates that while the effects of the disorders were somewhat consistent across datasets, ASD showed a relatively lower consistency between the datasets than MDD and SCZ. This suggests that the reason the FCs with higher effect size were not selected for ASD was the lower consistency of the disorder across datasets.

**Figure 7.**
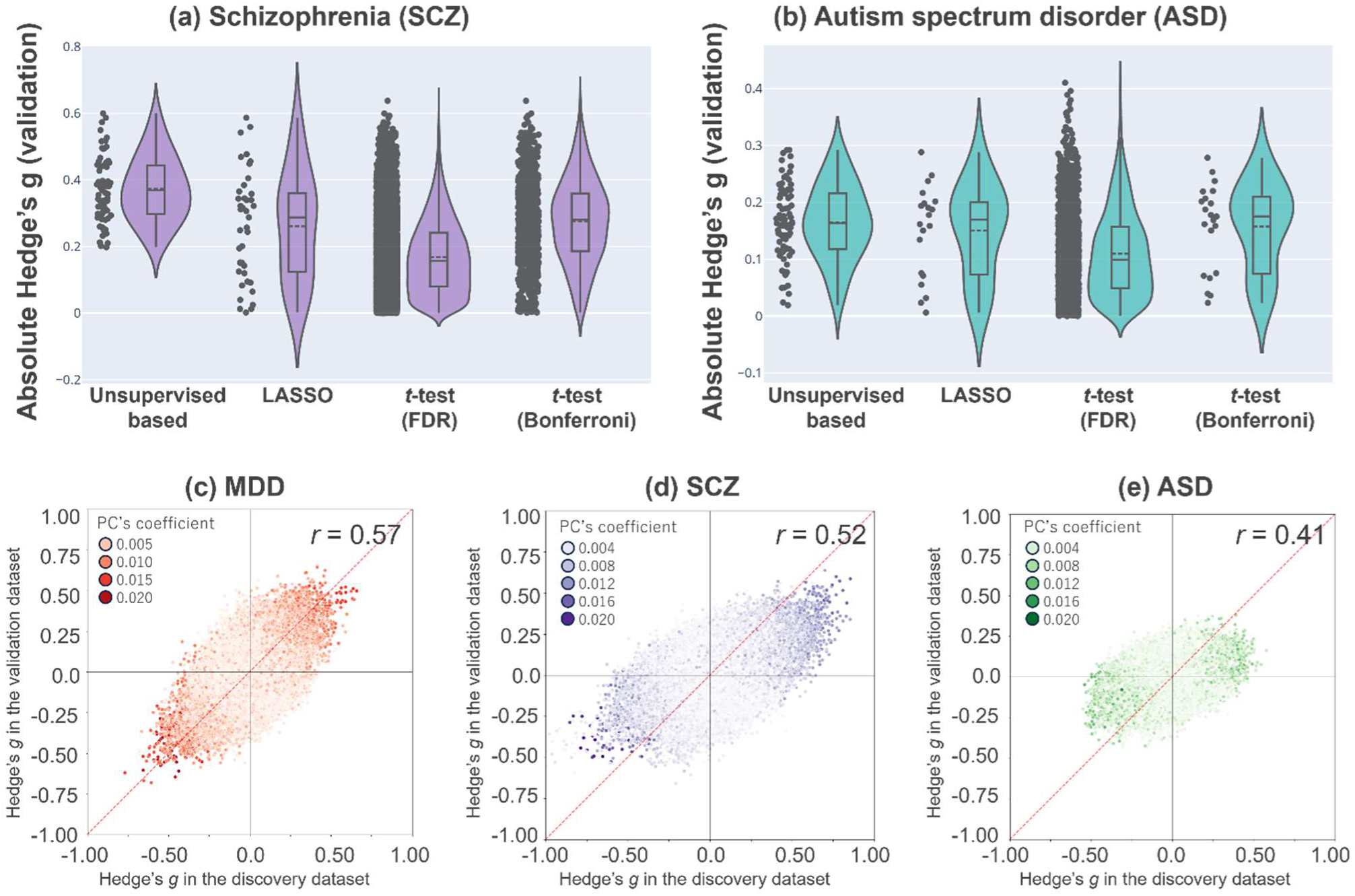
Validation in other psychiatric disorders. (a) Each gray point represents the effect size (Hedge’s *g*) of each FC for SCZ in the validation dataset, and the distribution is also shown. (b) Each gray point represents the effect size of each FC for ASD in the validation dataset, and the distribution is also shown. (c-e) Reproducibility across the two datasets (discovery and validation) regarding each diagnosis effect, respectively. Scatter plots of the diagnosis effect size are shown. The color of each point represents the weight of the PC related to each disorder, with darker colors indicating larger PC weight values. MDD: major depressive disorder, ASD, autism spectrum disorder, SCZ: schizophrenia, FC: functional connectivity, LASSO: Least Absolute Shrinkage and Selection Operator, FDR: false discovery rate

## 4. Discussion

In this study, we incorporated approximately 5,000 runs of rs-fMRI from six datasets, which included individuals with three different psychiatric disorders, and we demonstrated that an unsupervised feature computation-based feature selection method robustly extracted FCs related to psychiatric disorders compared to other conventional supervised feature selection methods. Furthermore, we found that the FCs selected by our proposed method had larger effect sizes than those selected by other conventional supervised feature selection methods.

The FCs selected by our proposed method were primarily involved in bilateral thalamic FCs, thalamo-motor FCs, and interhemispheric motor FCs in all three psychiatric disorders. The involvement of the thalamus in MDD, SCZ, and ASD has been discussed historically (Young et al. 2004; Long et al. 2024; Kim et al. 2017; Greicius et al. 2007; Chai et al. 2023; Müller et al. 2017; Myung et al. 2016; L.-N. Wang et al. 2024; Y. Wang et al. 2024; Segal et al. 2023; Kashiwagi et al. 2024; Northoff et al. 2021). Recent large-scale studies involving over 3,000 participants, including individuals with MDD, have consistently shown an increase in FC between the thalamus and cerebral cortex and a decrease in connectivity between bilateral cerebral cortical regions in individuals with MDD (Gallo et al. 2023), findings that align with those of our current study. Additionally, meta-analyses of effective connectivity, considering the directionality of FCs, have suggested that thalamic abnormalities serve as a hub in patients with MDD (Yang et al. 2023). At the neuronal level, an increase in the number of neurons in the thalamic region has been reported in postmortem brains of individuals with MDD (Young et al. 2004). While it was unclear whether there was an increase in excitatory projection neurons or inhibitory interneurons from the thalamus to the cerebral cortex, our study suggests an increase in connectivity between the thalamus and cerebral cortex, potentially indicating an increase in excitatory neurons in individuals with MDD. Furthermore, meta-analyses have reported reductions in thalamic volume (Boelens Keun et al. 2021) and alterations in microstructure (Zhang et al. 2022) in individuals with MDD. The interplay between FC changes, volumetric and microstructural alterations, and neuronal changes from macro to micro scales in the thalamus requires further investigation to elucidate the mechanisms linking these abnormalities to the symptoms of MDD.

Interestingly, it has been reported that arm immobilization for 2 weeks leads to decreased FC between the bilateral motor cortices and increased connectivity between the thalamus and motor cortex (Newbold et al. 2020; Chauvin et al. 2024). These changes are identical to those observed in patients with MDD in our current study. Considering that physical exercise is one of the effective treatments for MDD, its therapeutic effects may be mediated through the thalamus (Noetel et al. 2024). Additionally, a previous study reported that cognitive therapy reduces frontal-thalamic resting-state FC in social anxiety disorder (Kurita et al. 2023). To obtain more direct evidence, depressive symptoms were induced by deep brain stimulation applied to the thalamus in patients with Tourette Syndrome (Morishita et al. 2023). These results suggest that the thalamus plays a central role in depressive symptoms. Notably, since it is known that the thalamus consists of various subregions (B. Williams et al. 2024), future research will need to investigate which areas of the thalamus are more associated with MDD. It will also be important to examine how these subregions are related to specific symptoms of MDD.

Our proposed method selected FCs with the largest effect sizes not only in datasets of MDD but also in those of SCZ and ASD. However, in the ASD data, there was no significant difference in effect size compared to those of FCs selected by other methods. This could be due to the heterogeneity in ASD between the datasets. The inconsistency in findings regarding resting-state FC in datasets of ASD has already been discussed (He, Byrge, and Kennedy 2020), and our current analysis, which showed relatively low consistency of the ASD data between the discovery and validation datasets compared to the consistency of the data of other disorders, further supports this point. Moving forward, addressing the heterogeneity of diagnostic labels through initiatives, such as the Research Domain Criteria to redefine mental disorders, will be an important direction for research.

We found that utilizing these FCs did not enhance the predictive power of the brain network marker for psychiatric disorders. To investigate the reason, we examined the characteristics of the selected FCs using large-scale traveling subject data, which can separate various factors contributing to FC variance. The findings indicate that the selected FCs were susceptible to influences, such as individual differences, within-participant factors, imaging sites, and imaging protocols. Thus, while these FCs exhibited a large effect size in distinguishing psychiatric disorders, they are also prone to noise, which may compromise their utility as predictive features. This issue could be attributed to our use of PCA in the unsupervised method, as PCA does not differentiate whether variations originate from biological abnormalities or confounding factors. There are numerous dimension reduction techniques, and there is no guarantee that PCA is the optimal method or that it yields biologically valid dimensions. Determining the best unsupervised method remains a future challenge.

Despite the effectiveness of the proposed unsupervised feature computation-based feature selection method in identifying FCs with larger effect sizes associated with psychiatric disorders, several limitations should be noted. First, while it successfully extracted robust FCs distinguishing psychiatric disorders from healthy controls, it did not improve the predictive performance of the brain network marker, likely due to the influence of some nonbiological factors on the selected FCs. Second, the reliance on PCA for dimensionality reduction may not be optimal, as PCA identifies variance-maximizing components rather than biologically meaningful features. Third, although the proposed method reduced the reliance on diagnostic labels compared to conventional supervised approaches, it still used them. These limitations highlight the need for alternative dimensionality reduction techniques, strategies to address diagnostic label heterogeneity, and feature selection methods that are more robust to confounding factors in future research.

In this study that aimed to extract robust and generalizable FCs associated with psychiatric disorders from high-dimensional resting-state FC data, we proposed an unsupervised-based feature extraction method. Using our proposed method, we successfully identified key FCs involved in psychiatric disorders across various datasets. Interventions targeting these identified FCs could lead to effective treatments for psychiatric disorders.

## Supporting information

Supplementary Text S1: Harmonization method

Supplementary Text S2: Construction of the network marker and regularization-based feature selection method

Supplementary Text S3: FC variation analysis in the traveling-subject datasets

Supplementary Tables S1–4: Participants’ demographics in the datasets

Supplementary Tables S5–7: Imaging protocols used for the datasets

Supplementary Tables S8–S11: Selected FCs for MDD (discovery and validation), SCZ, and ASD

Supplementary Figure S1: Relationship between the principal component (PC) score and factors in the top 20 PCs in the schizophrenia (SCZ) discovery dataset

Supplementary Figure S2: Relationship between the principal component (PC) score and factors in the top 20 PCs in the autism spectrum disorder (ASD) discovery dataset

## Data available statement

Most data utilized in this study can be downloaded publicly from the DecNef Project Brain Data Repository from datasets (1) and (2) [https://bicr-resource.atr.jp/srpbsopen/], dataset (3) [https://bicr.atr.jp/dcn/en/download/harmonization/], dataset (5) [http://fcon_1000.projects.nitrc.org/indi/retro/cobre.html], and dataset (6) [https://fcon_1000.projects.nitrc.org/indi/abide/].

## Author Contributions

**Conceptualization**: Ayumu Yamashita, Okito Yamashita.

**Data curation**: Ayumu Yamashita, Takashi Itahashi, Masahiro Takamura, Hiroki Togo, Yujiro Yoshihara, Tomohisa Okada, Hirotaka Yamagata, Kenichiro Harada, Haruto Takagishi, Naohiro Okada, Osamu Abe, Takuya Hayashi, Shinsuke Koike, Saori C. Tanaka.

**Formal analysis**: Ayumu Yamashita.

**Funding acquisition**: Ayumu Yamashita, Yasumasa Okamoto, Go Okada, Hidehiko Takahashi, Kiyoto Kasai, Mitsuo Kawato, Okito Yamashita.

**Investigation**: Ayumu Yamashita, Takashi Itahashi, Yuki Sakai, Mitsuo Kawato, Hiroshi Imamizu, Okito Yamashita.

**Methodology**: Ayumu Yamashita, Yuki Sakai, Okito Yamashita.

**Project administration**: Ayumu Yamashita,Yasumasa Okamoto, Go Okada, Ryuichiro Hashimoto, Takashi Hanakawa, Toshiya Murai, Koji Matsuo, Hdehiko Takahashi, Kiyoto Kasai, Takuya Hayashi, Shinsuke Koike, Saori C. Tanaka, Mitsuo Kawato, Hiroshi Imamizu, Okito Yamashita.

**Software**: Ayumu Yamashita.

**Supervision**: Okito Yamashita.

**Visualization**: Ayumu Yamashita.

**Writing – original draft:** Ayumu Yamashita, Yuki Sakai, Mitsuo Kawato, Hiroshi Imamizu, Okito Yamashita.

**Writing – review & editing:** All authors

