## Supplementary Text for "Unsupervised feature computation-based feature selection robustly extracted resting-state functional connectivity patterns related to mental disorders"

**Supplementary Text S1: Travelling subject harmonization method**

We estimated the participant factor (***p***), measurement bias (***m***), sampling biases (***s***_hc_, ***s***_ssd_, ***s***_mdd_), and psychiatric disorder factor (***d***) by fitting the regression model to the functional connectivity (FC) values of all participants from both the discovery and travelling subject datasets, following the same method as in our previous study (A. Yamashita et al. 2019). As patients with autism spectrum disorder (ASD) were from only one site, we did not need to assume a sampling bias for ASD. For each FC, the regression model was formulated as follows:

$Connectivity={x_{m}}^{T}m+{x_{s_{hc}}}^{T}s_{hc}+{x_{s_{scz}}}^{T}s_{scz}+{{{x_{s_{mdd}}}^{T}s_{mdd}+x}_{d}}^{T}d+{x_{p}}^{T}p+const+\epsilon$ (Eq. S1)

$$such that\sum_{j}^{9} p_{j}=0,\sum_{k}^{4} m_{k}=0,\sum_{k}^{4} {s_{hc}}_{k}=0,\sum_{k}^{3} {s_{scz}}_{k}=0,{\sum_{k}^{3} {s_{mdd}}_{k}=0,d}_{1}\left( HC \right)=0$$

where $m$ is the measurement bias (4 sites × 1), $s_{hc}$ is the sampling bias for healthy controls (HCs) (4 sites × 1), $s_{scz}$ is the sampling bias for patients with schizophrenia (SCZ), $s_{mdd}$ is the sampling bias for patients with major depressive disorder (MDD), $d$ is the disorder factor (3 disorders × 1), $p$ is the participant factor (9 travelling subjects × 1), $const$ is the average FC value across all participants from all sites, and $\epsilon\sim\mathcal{N}\left( 0,\gamma^{-1} \right)$ denotes noise. A harmonized FC value was obtained by subtracting the estimated measurement bias from the following equation:

${Connectivity}^{Harmonised}=Connectivity-{x_{m}}^{t}\grave{m}$	(Eq. S2)

where $\grave{m}$ denotes the estimated measurement bias.

**Supplementary Text S2: Construction of the network marker and regularization-based feature selection**

### We constructed a brain network marker for MDD that distinguished between HCs and patients with MDD using the discovery dataset, based on 71,631 FC values, following the same method as in our previous study (A. Yamashita et al. 2020; Okada et al. 2023). To construct the network marker, we applied a machine learning technique called logistic regression with Least Absolute Shrinkage and Selection Operator (LASSO) regularization, as we assumed that psychiatric disorder factors were associated with a specific subset of connections. A logistic function was used to define the probability of a participant belonging to the MDD class as follows:

$P_{sub}\left( y_{sub}=1|\boldsymbol{c}_{sub};\boldsymbol{w} \right)=\frac{1}{1+\exp\left( -\boldsymbol{w}^{T}\boldsymbol{c}_{sub} \right)}$, (Eq. S3)

### in which $\boldsymbol{y}_{\boldsymbol{sub}}$ $\mathbf{y}_{\mathbf{sub}}$represents the class label (MDD, *y* = 1; HC, *y* = 0) of a participant, $\mathbf{c}_{\mathbf{sub}}$ $\boldsymbol{c}_{\boldsymbol{sub}}$ represents an FC vector for a given participant, and *w* represents the weight vector. The weight vector *w* was determined to minimize

$J\left( \mathbf{w} \right)=-\frac{1}{n_{sub}}\sum_{j=1}^{n_{sub}} \log P_{j}\left( y_{j}=1|\boldsymbol{c}_{\boldsymbol{j}};\boldsymbol{w} \right)+\lambda\left\| \boldsymbol{w} \right\|_{1},$ (Eq. S4)

### in which $\left\| \boldsymbol{w} \right\|_{\boldsymbol{1}}\boldsymbol{=}\sum_{\boldsymbol{i}}^{\boldsymbol{N}} \left| \boldsymbol{w}_{\boldsymbol{i}} \right|$ and $\boldsymbol{\lambda}$ represent hyperparameters that control the amount of shrinkage applied to the estimates. To determine the weights of the logistic regression model and hyperparameter λ, we employed a nested cross-validation (CV) approach. In this process, the discovery dataset was first split into a training set (9 out of 10 folds) for model training and test set (1 out of 10 folds) for evaluation. To address potential biases caused by the imbalance between the number of patients with MDD and HCs, we applied an undersampling technique. Specifically, we randomly selected approximately 125 MDD patients and 125 HCs from the training set, and the classifier’s performance was assessed using the test set. Furthermore, to ensure comparable age distributions between the MDD and HC groups within each subsample, we adjusted the mean age during the undersampling and subsampling steps. Given that only a subset of the training data was used after undersampling, we repeated this random sampling procedure 10 times (subsampling). For each subsample, a model was trained while tuning the regularization parameter within the inner loop of the nested CV, ultimately producing 10 classifiers.

For the inner loop, we used the “*lassoglm*” function in MATLAB (R2016b, Mathworks, USA) with parameters set to "NumLambda = 25" and "CV = 10". The regularization path was defined by first determining the maximum $\lambda$ ($\lambda_{\max}$), which ensured that the optimal solution was an all-zero vector. A total of 25 $\lambda$ values were then selected at equal intervals from 0 to $\lambda_{\max}$. The final$\lambda$was chosen using the one-standard-error rule, which selects the largest λ within one standard deviation of the minimum prediction error. The classifier output (diagnostic probability) was averaged across the 10 models, and individuals with a diagnostic probability > 0.5 were classified as MDD patients. The area under the curve (AUC) was computed using the "perfcurve" function in MATLAB. In addition, we evaluated classification performance using the accuracy, sensitivity, specificity, positive predictive value (PPV), and negative predictive value (NPV). The Matthews correlation coefficient (MCC) was also calculated to assess model performance in imbalanced datasets. To evaluate the generalizability of the brain network marker, we applied the trained classifiers to an independent validation dataset. Since 100 classifiers were generated through 10-fold CV × 10 subsamplings, all trained models were used to classify the validation dataset. The final diagnostic probability for each participant was obtained by averaging the outputs of the 100 classifiers, and individuals with a probability > 0.5 were classified as MDD patients.

### To identify the most diagnostically relevant FCs, we examined the frequency with which each FC was selected by LASSO during the 10-fold CV process. An FC was considered important if its selection frequency exceeded the chance level, as determined by a permutation test (regularization-based feature selection). Specifically, diagnostic labels in the discovery dataset were permuted, and the entire 10-fold CV and 10-subsampling procedure was repeated 100 times. The selection count for each connection across 10-fold CV × 10 subsamplings (maximum 100 times) was used as a statistic for each permutation dataset. To control for multiple comparisons, we established a null distribution by taking the maximum selection count across all functional connections and set the statistical significance threshold at P < 0.05 (one-sided). FCs selected at least 17 times out of 100 were considered diagnostically significant.

**Supplementary Text S3: FC variation analysis in the traveling subject datasets**

To assess the impact of experimental factors, such as participants, scanners, and imaging protocols, on FC, we applied a linear fixed-effects model to each FC, following the same method as in our previous study (O. Yamashita et al. 2024). This approach enabled us to estimate the extent to which these factors influenced FC. Specifically, we used a three-factor model—including participant, scanner, and imaging protocol—when analyzing the BMB traveling subject dataset, while for the SRPBS traveling subject dataset, we employed a two-factor model consisting of participant and scanner effects.

$z_{nc}$ represents the z-transformed FC strength for a given run 𝑛 and a specific FC 𝑐, where 𝑁 is the total number of runs and 𝐶 is the total number of FCs. We defined $\boldsymbol{z}_{c}=\left( z_{1c}, z_{2c},\ldots, z_{nc} \right)$ as a column vector containing the FC strengths across all runs for a particular FC 𝑐. In the case of the three-factor model, the relationship was modeled using a linear regression equation with three explanatory variables:

$$\boldsymbol{z}_{c}=X_{p}\boldsymbol{\beta}_{\boldsymbol{c}}^{\boldsymbol{p}}\boldsymbol{+}X_{prot}\boldsymbol{\beta}_{\boldsymbol{c}}^{\boldsymbol{prot}}\boldsymbol{+}X_{scan}\boldsymbol{\beta}_{\boldsymbol{c}}^{\boldsymbol{scan}}\boldsymbol{+}\boldsymbol{\epsilon}_{\boldsymbol{c}}\boldsymbol{,}(Eq. S5)$$

Here, the three factors were treated as categorical variables, represented by binary matrices $X_{p}, X_{prot},$and $X_{scan}$. For instance, the participant-factor matrix $X_{p}$was structured as an *N* × *P* matrix (where *P* is the total number of participants), with its (*i*, *j*) element set to 1 if participant 𝑖 was involved in run *j*, and otherwise it was to 0. The parameter vector $\boldsymbol{\beta}_{\boldsymbol{c}}^{\boldsymbol{p}}$**,** with dimensions *P* × 1, captured the corresponding effect magnitude for each participant. Similar definitions were applied to the protocol and scanner factors. The term $\boldsymbol{\epsilon}_{\boldsymbol{c}}$represented residuals that could not be accounted for by the linear combination of these three factors.

The model (Eq. S5) can be rewritten in a more compact form:

$$\boldsymbol{z}_{c}=\boldsymbol{X}\boldsymbol{\beta}_{c}\boldsymbol{+}\boldsymbol{\epsilon}_{\boldsymbol{c}}\boldsymbol{,}(Eq. S6)$$

where $\boldsymbol{X}\boldsymbol{=}\left[ X_{p} X_{prot} X_{scan} \right]$ and $\boldsymbol{\beta}_{c}= \left[ {\boldsymbol{\beta}_{\boldsymbol{c}}^{\boldsymbol{p}}}^{\boldsymbol{t}} {\boldsymbol{\beta}_{\boldsymbol{c}}^{\boldsymbol{prot}}}^{\boldsymbol{t}} {\boldsymbol{\beta}_{\boldsymbol{c}}^{\boldsymbol{scan}}}^{\boldsymbol{t}} \right]^{t}$ are the combined explanatory matrix and parameter vector, respectively. Using the least squares method, the parameter vector $\boldsymbol{\beta}_{c}$was estimated by solving the normal equation:

$$\boldsymbol{X}^{t}\boldsymbol{X}\boldsymbol{\beta}_{c}=\boldsymbol{X}^{t}\boldsymbol{z}_{c}, (Eq. S7)$$

Since all three factors were categorical, the matrix $\boldsymbol{X}$ was not of full rank. This could be verified by summing the columns of $X_{p}, X_{prot},$and $X_{scan}$, which would result in a vector of ones. Consequently, the linear equation ($Eq. S7$) lacked a unique solution (and the inverse of $\boldsymbol{X}^{t}\boldsymbol{X}$ does not exist). Instead, the least squares solution was obtained using the Moore-Penrose pseudo-inverse:

$$\boldsymbol{\beta}_{c}=\left( \boldsymbol{X}^{t}\boldsymbol{X} \right)^{\dagger}\boldsymbol{X}^{t}\boldsymbol{z}_{c}, (Eq. S8)$$

This solution minimized the L2-norm. Based on statistical analysis, the baseline values for each factor were arbitrary, and only the relative differences within each factor had meaningful interpretability. Therefore, we computed the FC variations attributed to the participant (or individual subject), protocol, and scanner factors by determining the standard deviation of $\boldsymbol{\beta}_{\boldsymbol{c}}^{\boldsymbol{p}}\boldsymbol{,}\boldsymbol{\beta}_{\boldsymbol{c}}^{\boldsymbol{prot}},$and $\boldsymbol{\beta}_{\boldsymbol{c}}^{\boldsymbol{scan}}$ across the members within each factor, respectively. The pair-wise distance matrix between members of the scanner type and imaging protocol factors was computed by calculating the mean absolute difference between corresponding estimated parameters across all FCs.

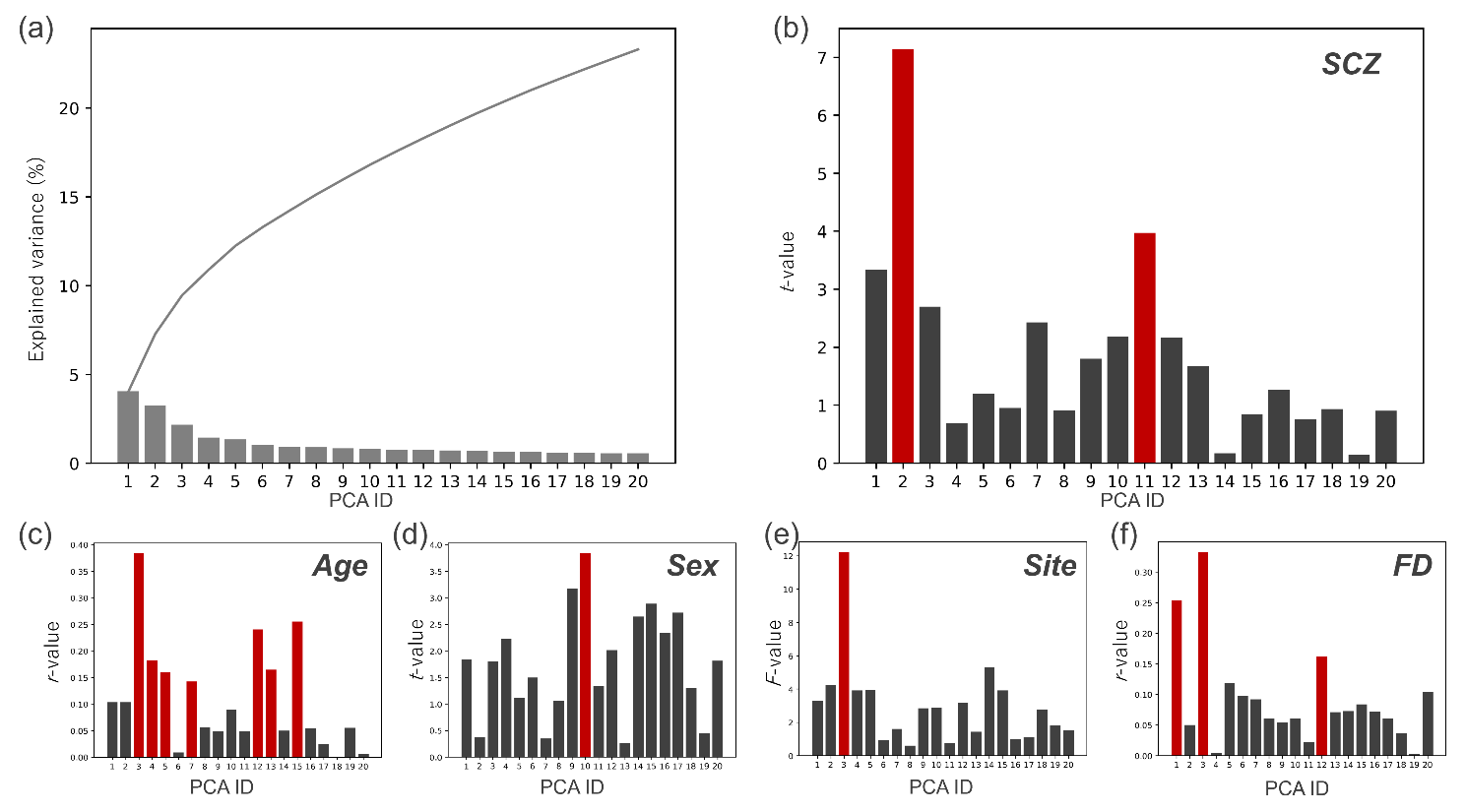

**Supplementary Figure S1.** **Relationship between the principal component (PC) score and factors in the top 20 PCs in the schizophrenia (SCZ) discovery dataset:** (a) Explained variance of the top 20 PCs and their cumulative summation. (b–f) Relationship between each factor and the top 20 PCs. (b) Difference in PC scores between HCs and patients with SCZ (*t*-value). (c) Pearson’s correlation coefficients between the PC score and age. (d) Difference in PC scores between men and women (*t*-value). (e) Difference in PC scores across imaging sites (*F*-value). (f) Pearson’s correlation coefficients between the PC score and head motion (framewise displacement value). The red bar shows a significant relationship with the factor. Here, we only visualized the top 20 PCs for visualization purpose. Of note, we used all PCs for the analyses.

PCA: principal component analysis

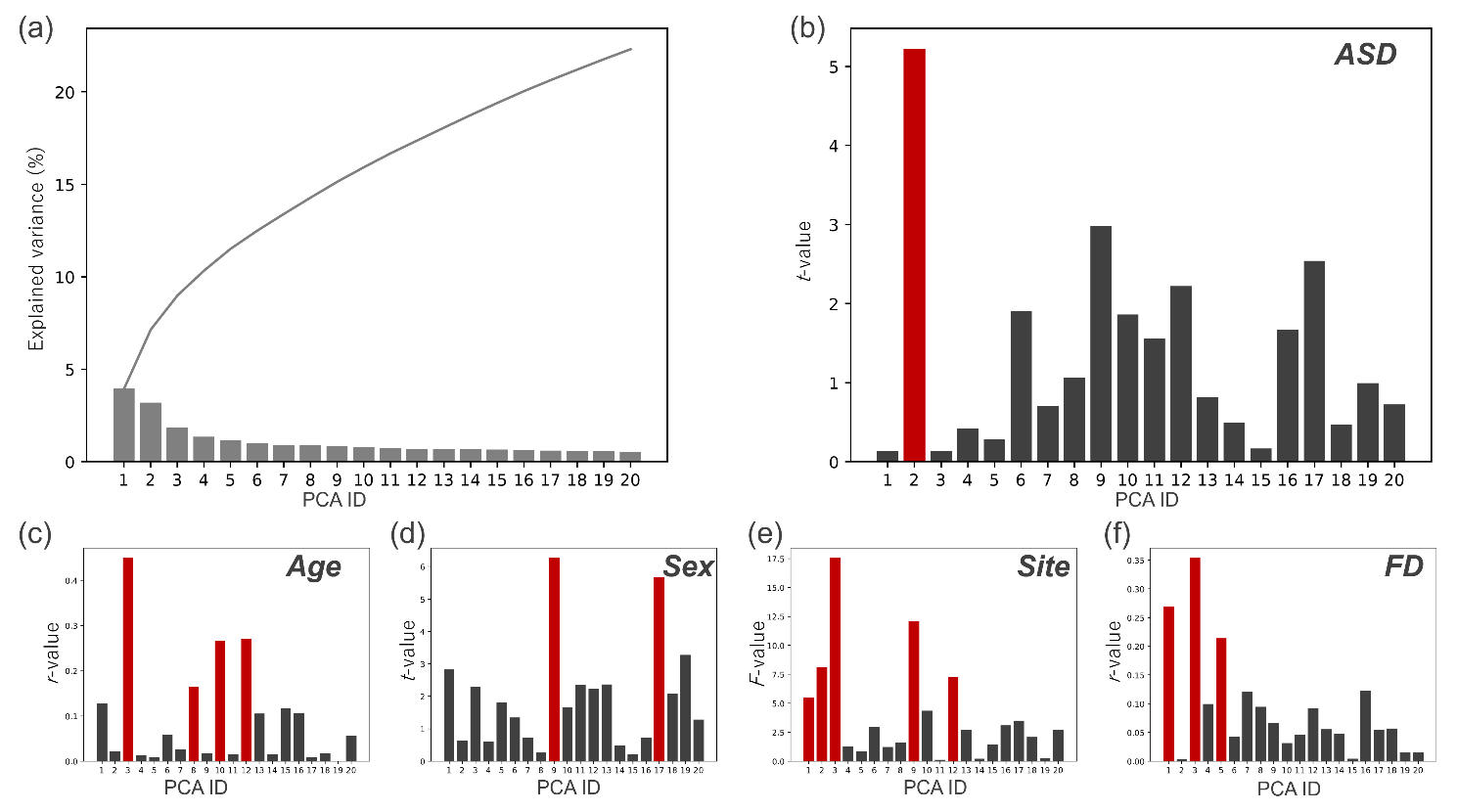

**Supplementary Figure S2.** **Relationship between the principal component (PC) score and factors in the top 20 PCs in the autism (ASD) discovery dataset:** (a) Explained variance of the top 20 PCs and their cumulative summation. (b–f) Relationship between each factor and the top 20 PCs. (b) Difference in PC scores between HCs and patients with ASD (*t*-value). (c) Pearson’s correlation coefficients between the PC score and age. (d) Difference in PC scores between men and women (*t*-value). (e) Difference in PC scores across imaging sites (*F*-value). (f) Pearson’s correlation coefficients between the PC score and head motion (framewise displacement value). The red bar shows a significant relationship with the factor. Here, we only visualized the top 20 PCs for visualization purpose. Of note, we used all PCs for the analyses.
