## Supplementary Tables for "Unsupervised feature computation-based feature selection robustly extracted resting-state functional connectivity patterns related to mental disorders"

**Supplementary Table S1. Participant demographics in the “discovery dataset”**

| **Site** | **HC** | | | **MDD** | | | **ASD** | | | | **SCZ** | | | **ALL** | | |
| --- | --- | --- | --- | --- | --- | --- | --- | --- | --- | --- | --- | --- | --- | --- | --- | --- |
|  | N | M/F | Age (mean±SD) | N | M/F | Age (mean±SD) | N | M/F | Age (mean±SD) | N | | M/F | Age (mean±SD) | N | M/F | Age (mean±SD) |
| SWA | 99 | 84/15 | 28.4±7.9 | – | – | – | 111 | 96/15 | 32.0±7.5 | 18 | | 14/4 | 42.8±8.6 | 228 | 194/34 | 31.3±8.6 |
| COI | 112 | 44/68 | 50.9±13.4 | 62 | 29/33 | 44.6±12.4 | – | – | – | – | | – | – | 174 | 73/101 | 48.7±13.4 |
| KUT | 166 | 99/67 | 35.6±13.5 | 17 | 11/6 | 43.9±13.3 | – | – | – | 48 | | 24/24 | 41.5±10.4 | 231 | 134/97 | 37.5±13.2 |
| UTO | 168 | 77/91 | 35.6±17.5 | 59 | 35/24 | 38.2±11.4 | 10 | 9/1 | 37.0±9.6 | 36 | | 24/12 | 31.4±10.3 | 273 | 145/128 | 35.7±15.4 |
| Summary | 545 | 304/241 | 37.4±15.9 | 138 | 75/63 | 41.8±12.4 | 121 | 105/16 | 32.4±7.8 | 102 | | 62/40 | 38.2±11.2 | 906 | 546/360 | 37.5±14.3 |
| **Abbreviations**: HC: healthy control, MDD: major depressive disorder, ASD: autism spectrum disorder, SCZ: schizophrenia, M: Male, F: Female, SD: standard deviation. SWA: Showa University, COI: Centre of Innovation, Hiroshima University, KUT: Kyoto University (TimTrio), UTO: the University of Tokyo | | | | | | | | | | | | | | | | |

**Supplementary Table S2. Participant demographics in the “validation dataset”**

| **Site** | **HC** | | | **MDD** | | | **ASD** | | | **SCZ** | | | **ALL** | | | |
| --- | --- | --- | --- | --- | --- | --- | --- | --- | --- | --- | --- | --- | --- | --- | --- | --- |
|  | N | M/F | Age (mean±SD) | N | M/F | Age (mean±SD) | N | M/F | Age (mean±SD) | N | M/F | Age (mean±SD) | | N | M/F | Age (mean±SD) |
| HUH | 66 | 29/37 | 34.6±13.0 | 57 | 32/25 | 43.3±12.2 | – | – | – | – | – | – | | 123 | 61/62 | 38.6±13.3 |
| HRC | 49 | 13/36 | 41.7±11.7 | 16 | 6/10 | 40.5±11.5 | – | – | – | – | – | – | | 65 | 19/46 | 41.4±11.5 |
| HKH | 28 | 11/17 | 45.9±9.3 | 32 | 20/12 | 45.2±11.5 | – | – | – | – | – | – | | 60 | 31/29 | 45.5±10.5 |
| KTT | 75 | 48/27 | 28.9±9.1 | – | – | – | – | – | – | 52 | 27/25 | 37.2±9.4 | | 127 | 75/52 | 32.3±10.0 |
| UYA | 120 | 50/70 | 45.9±19.5 | 76 | 35/41 | 50.2±13.8 | – | – | – | – | – | – | | 196 | 85/111 | 47.6±17.6 |
| Summary | 338 | 151/187 | 39.3±16.1 | 181 | 93/88 | 46.3±13.1 | – | – | – | 52 | 27/25 | 37.2±9.4 | | 571 | 271/300 | 41.3±15.1 |
| **Abbreviations**: HC: healthy control, SCZ: schizophrenia, MDD: major depressive disorder, ASD: autism spectrum disorder, M: Male, F: Female, SD: standard deviation. HUH: Hiroshima University Hospital, HRC: Hiroshima Rehabilitation Center, HKH: Hiroshima Kajikawa Hospital, KTT: Kyoto University (Trio), UYA: Yamaguchi University | | | | | | | | | | | | | | | | |

**Supplementary Table S3. Participant demographics in the “COBRE dataset”**

| **Site** | **HC** | | | | **MDD** | | | **ASD** | | | **SCZ** | | | | **ALL** | | | |
| --- | --- | --- | --- | --- | --- | --- | --- | --- | --- | --- | --- | --- | --- | --- | --- | --- | --- | --- |
|  | N | M/F | Age (mean±SD) | N | | M/F | Age (mean±SD) | N | M/F | Age (mean±SD) | | N | M/F | Age (mean±SD) | | N | M/F | Age (mean±SD) |
| COBRE | 45 | 31/14 | 32.0±9.6 | – | | – | – | – | – | – | | 30 | 26/4 | 31.8±12.2 | | 75 | 57/18 | 31.9±10.6 |
| **Abbreviations**: HC: healthy control, MDD: major depressive disorder, ASD: autism spectrum disorder, SCZ: schizophrenia, M: Male, F: Female, SD: standard deviation. COBRE: Centre of Biomedical Research Excellence | | | | | | | | | | | | | | | | | | |

**Supplementary Table S4. Participant demographics in the “ABIDE-I and -II dataset”**

|  | **HC** | | | **MDD** | | | | | **ASD** | | | **SCZ** | | | | **ALL** | | |
| --- | --- | --- | --- | --- | --- | --- | --- | --- | --- | --- | --- | --- | --- | --- | --- | --- | --- | --- |
|  | N | M/F | Age (mean±SD) | | N | M/F | Age (mean±SD) | N | | M/F | Age (mean±SD) | N | M/F | Age (mean±SD) | N | | M/F | Age (mean±SD) |
| CMU_a | 8 | 7/1 | 26.4±4.3 | | – | – | – | 6 | | 6/0 | 26.2±4.7 | – | – | – | 14 | | 13/1 | 26.3±4.3 |
| CMU_b | 5 | 3/2 | 27.6±8.1 | | – | – | – | 7 | | 5/2 | 25.9±7.2 | – | – | – | 12 | | 8/4 | 26.6±7.3 |
| KKI | 32 | 23/9 | 10.2±1.3 | | – | – | – | 14 | | 11/3 | 9.8±1.4 | – | – | – | 46 | | 34/12 | 10.0±1.3 |
| Leuven_1 | 15 | 15/0 | 23.3±2.9 | | – | – | – | 14 | | 14/0 | 21.9±4.1 | – | – | – | 29 | | 29/0 | 22.6±3.6 |
| Leuven_2 | 20 | 15/5 | 14.3±1.5 | | – | – | – | 14 | | 11/3 | 13.7±1.1 | – | – | – | 34 | | 26/8 | 14.1±1.4 |
| MaxMun | 12 | 8/4 | 33.2±9.0 | | – | – | – | 12 | | 9/3 | 35.5±11.4 | – | – | – | 24 | | 17/7 | 34.4±10.1 |
| MaxMun2 | 18 | 18/0 | 23.0±7.5 | | – | – | – | 9 | | 9/0 | 19.2±12.8 | – | – | – | 27 | | 27/0 | 21.7±9.5 |
| NYU | 104 | 78/26 | 15.8±6.3 | | – | – | – | 76 | | 66/10 | 14.8±7.0 | – | – | – | 180 | | 144/36 | 15.4±6.6 |
| Olin | 12 | 10/2 | 17.8±3.2 | | – | – | – | 13 | | 11/2 | 17.2±3.7 | – | – | – | 25 | | 21/4 | 17.5±3.4 |
| Pitt | 22 | 19/3 | 19.1±6.4 | | – | – | – | 19 | | 16/3 | 19.3±7.2 | – | – | – | 41 | | 35/6 | 19.2±6.7 |
| SBL | 15 | 15/0 | 33.7±6.6 | | – | – | – | 14 | | 14/0 | 35.3±10.8 | – | – | – | 29 | | 29/0 | 34.5±8.7 |
| Trinity | 23 | 23/0 | 17.5±3.7 | | – | – | – | 21 | | 21/0 | 17.5±3.2 | – | – | – | 44 | | 44/0 | 17.5±3.4 |
| UCLA_1 | 28 | 24/4 | 13.6±2.0 | | – | – | – | 29 | | 25/4 | 13.6±2.7 | – | – | – | 57 | | 49/8 | 13.6±2.3 |
| UCLA_2 | 12 | 10/2 | 12.4±1.0 | | – | – | – | 8 | | 8/0 | 12.4±2.1 | – | – | – | 20 | | 18/2 | 12.4±1.5 |
| UM | 75 | 57/18 | 14.8±3.6 | | – | – | – | 54 | | 45/9 | 13.7±2.3 | – | – | – | 129 | | 102/27 | 14.3±3.2 |
| USM | 41 | 41/0 | 21.7±7.5 | | – | – | – | 51 | | 51/0 | 23.1±7.9 | – | – | – | 92 | | 92/0 | 22.5±7.7 |
| Yale | 26 | 19/7 | 12.8±2.8 | | – | – | – | 26 | | 19/7 | 12.8±3.0 | – | – | – | 52 | | 38/14 | 12.8±2.9 |
| ABIDEII  BNI_1 | 23 | 23/0 | 40.0±15.7 | | – | – | – | 24 | | 24/0 | 37.1±15.8 | – | – | – | 47 | | 47/0 | 38.5±15.6 |
| ABIDEII  EMC_1 | 15 | 12/3 | 8.1±1.0 | | – | – | – | 14 | | 12/2 | 8.6±1.3 | – | – | – | 29 | | 24/5 | 8.3±1.1 |
| ABIDEII  GU_1 | 30 | 16/14 | 10.8±1.8 | | – | – | – | 20 | | 18/2 | 11.4±1.6 | – | – | – | 50 | | 34/16 | 11.0±1.8 |
| ABIDEII  IU_1 | – | – | – | | – | – | – | 2 | | 1/1 | 31.5±14.8 | – | – | – | 2 | | 1/1 | 31.5±14.8 |
| ABIDEII  KKI_8 | 186 | 108/78 | 10.4±1.2 | | – | – | – | 56 | | 44/12 | 10.6±1.5 | – | – | – | 242 | | 152/90 | 10.4±1.3 |
| ABIDEII  KKI_32 | 78 | 56/22 | 10.5±1.2 | | – | – | – | 20 | | 16/4 | 10.8±1.4 | – | – | – | 98 | | 72/26 | 10.5±1.3 |
| ABIDEII  KUL_3 | – | – | – | | – | – | – | 2 | | 2/0 | 20.5±0.7 | – | – | – | 2 | | 2/0 | 20.5±0.7 |
| ABIDEII  NYU_1 | 28 | 26/2 | 9.5±3.4 | | – | – | – | 43 | | 39/4 | 10.2±6.0 | – | – | – | 71 | | 65/6 | 9.9±5.1 |
| ABIDEII  OHSU_1 | 50 | 26/24 | 10.5±1.7 | | – | – | – | 31 | | 25/6 | 11.6±2.3 | – | – | – | 81 | | 51/30 | 10.9±2.0 |
| ABIDEII  ONRC_2 | 32 | 19/13 | 24.4±3.5 | | – | – | – | 20 | | 17/3 | 21.8±3.5 | – | – | – | 52 | | 36/16 | 23.4±3.7 |
| ABIDEII  TCD_1 | 18 | 18/0 | 16.3±2.8 | | – | – | – | 14 | | 14/0 | 15.3±3.5 | – | – | – | 32 | | 32/0 | 15.9±3.1 |
| ABIDEII  UCD_1 | 13 | 9/4 | 15.0±1.6 | | – | – | – | 15 | | 12/3 | 15.0±2.0 | – | – | – | 28 | | 21/7 | 15.0±1.8 |
| ABIDEII  UCLA_1 | 13 | 9/4 | 10.0±2.3 | | – | – | – | 11 | | 11/0 | 12.2±1.7 | – | – | – | 24 | | 20/4 | 11.0±2.3 |
| ABIDEII  USM_1 | 15 | 12/3 | 24.0±8.1 | | – | – | – | 15 | | 13/2 | 19.2±6.9 | – | – | – | 30 | | 25/5 | 21.6±7.8 |
| Summary | 969 | 719/250 | 15.3±8.1 | | – | – | – | 674 | | 589/85 | 16.5±9.1 | – | – | – | 1643 | | 1308/335 | 15.8±8.6 |
| **Abbreviations**: ABIDE: Autism Brain Imaging Data Exchange, HC: healthy control, MDD: major depressive disorder, ASD: autism spectrum disorder, SCZ: schizophrenia, M: Male, F: Female, SD: standard deviation. CMU: Carnegie Mellon University, KKI: Kennedy Krieger Institute, MaxMun: Ludwig Maximilians University Munich, NYU: New York University Langone Medical Center, SBL: Social Brain Lab BCN NeuroImaging Center, University Medical Center Groningen and Netherlands Institute for Neurosciences, UCLA: the University of California, Los Angeles, University of Michigan, USM: University of Utah School of Medicine, BNI: Barrow Neurological Institute, EMC: Erasmus University Medical Center Rotterdam, GU: Georgetown University, IU: Indiana University, KUL: Katholieke Universiteit Leuven, OHSU: Oregon Health and Science University, ONRC: Olin Neuropsychiatry Research Center, Institute of Living at Hartford Hospital, TCD: Trinity Centre for Health Sciences, UCD: University of California Davis, | | | | | | | | | | | | | | | | | | |

**Supplementary Table S5. Imaging parameters of each imaging site in the discovery, validation and COBRE datasets.**

|  | **Discovery dataset** | | | | **Validation dataset** | | | | | | | **COBRE** |
| --- | --- | --- | --- | --- | --- | --- | --- | --- | --- | --- | --- | --- |
| Site | KUT | SWA | COI | UTO | KTT | KUP | HKH | | HRC | | HUH | COBRE |
| MRI scanner | Siemens TimTrio | Siemens Verio | Siemens Verio | GE MR750w | Siemens Trio | Siemens Prisma | Siemens Spectra | | GE Signa HDxt | | GE Signa HDxt | Siemens TimTrio |
| Magnetic field strength | 3.0 T | | | | | | | | | | | |
| Channels per coil | 32 | 12 | | 24 | 8 | 64 | 12 | | 8 | | | |
| Field of view (mm) | 212 × 212 | | | | 256 × 192 | 200 × 200 | 192 × 192 | | 256 × 256 | | | 240 × 240 |
| Matrix | 64 × 64 | | | | 64 × 48 | 100 × 100 | 64 × 64 | | | | | |
| Number of slices | 40 | | | | 30 | 72 | 38 | | 32 | | | 33 |
| Number of volumes | 240 | | | | 180 | 320 | 107 | | 143 | | | 150 |
| In-plane resolution (mm) | 3.3125 × 3.3125 | | | | 4 × 4 | 2 × 2 | 3 × 3 | | 4 × 4 | | | 3.75 × 3.75 |
| Slice thickness (mm) | 3.2 | | | | 4 | 2 | 3 | | 4 | | | 3.5 |
| Slice gap (mm) | 0.8 | | | | 0 | | | | | | | 1.05 |
| TR (s) | 2.5 | | | | 2 | 0.75 | 2.7 | | 2 | | | |
| TE (ms) | 30 | | | | 30 | 36.2 | 31 | | 27 | | | 29 |
| Total scan time | 10'00" | | | | 6'00" | 4'10" | 5'00" | | 4'46" | | 5'00" | 6'00" |
| Flip angle (degree) | 80 | | | | 90 | 55 | 90 | | | | | 75 |
| Slice acquisition order | Ascending | | | | Ascending (interleaved) | | | Ascending | | Ascending (interleaved) | | |
| Phase encoding | P→A | | A→P | P→A | A→P | P→A | A→P | | | | P→A | A→P |
| Eye condition | Fixated | | | | Fixated | | | | | | | Not specified |

**Supplementary Table S6. Imaging parameters of each imaging site in the ABIDE-I dataset.**

| Site | CMU | KKI | Leuven | MaxMun | NYU | Olin | Pitt | SBL | Trinity | UCLA | UM | USM | Yale |
| --- | --- | --- | --- | --- | --- | --- | --- | --- | --- | --- | --- | --- | --- |
| MRI scanner | Siemens  Verio | Philips  Achieva | Philips  Intera | Siemens  Verio | Siemens  Allegra | Siemens  Allegra | Siemens  Allegra | Philips Intera | Philips  Achieva | Siemens  TrioTim | GE  Signa | Siemens  TrioTim | Siemens  TimTrio |
| Magnetic field strength | 3T | 3T | 3T | 3T | 3T | 3T | 3T | 3T | 3T | 3T | 3T | 3T | 3T |
| Channels per coil | NA | 8 | 8 | NA | NA | NA | NA | NA | 8 | NA | NA | NA | NA |
| Field of view (mm) | 192x192 | 256x256 | 230x230 | 192x192 | 240x192 | 220x220 | 200x200 | 220x220 | 240x240 | 192x192 | 220x220 | 220x220 | 220x220 |
| Matrix | 64x64 | 84x81 | 64x64 | 64x64 | 80x64 | 64x64 | 64x64 | 80x80 | 80x80 | 64x64 | 64x64 | 64x64 | 64x64 |
| Number of slices | 21 or 28 | 47 | 32 | 28 | 33 | 29 | 29 | 38 | 38 | 34 | 40 | 40 | 34 |
| Number of volumes | 240 | 156 | 250 | 120 | 180 | 210 | 200 | 200 | 150 | 120 | 300 | 240 | 200 |
| In-plane resolution (mm) | 3x3 | 3.05x3.15 | 3.59x3.59 | 3x3 | 3x3 | 3.4x3.4 | 3.1x3.1 | 2.75x2.75 | 3x3 | 3x3 | 3.438x3.438 | 3.4x3.4 | 3.4x3.4 |
| Slice thickness (mm) | 3 | 3 | 4 | 4 | 4 | 4 | 4 | 2.72 | 3.5 | 4 | 3 | 3 | 4 |
| Slice gap (mm) | 1.5 | 0 | 0 | 0.4 | 0 | 1 | 0 | 0.272 | 0.35 | 0 | 0 | 0.3 | 0 |
| TR (s) | 1.5 or 2 | 2.5 | 1.667 | 3 | 2 | 1.5 | 1.5 | 2.2 | 2 | 3 | 2 | 2 | 2 |
| TE (ms) | 0.03 | 0.03 | 0.033 | 0.03 | 0.015 | 0.027 | 0.025 | 0.03 | 0.028 | 0.028 | 0.03 | 0.028 | 0.025 |
| Total scan time | 8:06 | 6:40 | 7:06 | 6:06 | 6:00 | 5:15 | 5:06 | 7:28 | 5:06 | 6:06 | 10:00 | 8:06 | 6:40 |
| Flip angle (degree) | 73 | 75 | 90 | 80 | 90 | 60 | 70 | 80 | 90 | 90 | 90 | 90 | 60 |
| Slice acquisition order | Ascending (interleaved) | Ascending (sequential) | Ascending (sequential) | Ascending (interleaved) | Ascending (interleaved) | Ascending (interleaved) | Ascending (interleaved) | Descending  (sequential) | Ascending (sequential) | Ascending (interleaved) | Ascending (sequential) | Ascending (interleaved) | Ascending (interleaved) |
| Phase encoding | AP | AP | AP | AP | RL | AP | AP | AP | AP | AP | AP | AP | AP |
| Eye condition | Close | Fixated | Close | Close or Open | Fixated | Fixated | Close | Close | Close | Open | Fixated | Open | Open |

**Supplementary Table S7. Imaging parameters of each imaging site in the ABIDE-II dataset.**

| Site | BNI | EMC | GU | IU | KKI | KUL | NYU | OHSU | ONRC | TCD | UCD | UCLA | USM |
| --- | --- | --- | --- | --- | --- | --- | --- | --- | --- | --- | --- | --- | --- |
| MRI scanner | Philips  Ingenia | GE  MR750 | Siemens  TrioTim | Siemens  TrioTim | Philips  Achieva | Philips  Achieva | Siemens  Allegra | Siemens  TrioTim | Siemens  Skyra | Philips Intera Achieva | Siemens  TrioTim | Siemens  TrioTim | Siemens  TrioTim |
| Magnetic field strength | 3T | 3T | 3T | 3T | 3T | 3T | 3T | 3T | 3T | 3T | 3T | 3T | 3T |
| Channels per coil | 15 | 8 | 12 | 32 | 8 or 32 | 32 | 8 | 12 | NA | NA | 32 | NA | 12 |
| Field of view (mm) | 240X240 | 230x230 | 192x192 | 220x220 | 256x256 | 200x200 | 240x240 | 240x240 | 240x240 | 240x240 | 224x224 | 192x192 | 220x220 |
| Matrix | 64X64 | 64x64 | 64x64 | 64x64 | 84x81 | 80x78 | 80x80 | 64x64 | 80x80 | 80x80 | 64x64 | 64x64 | 64x64 |
| Number of slices | 50 | 37 | 43 | 42 | 47 | 45 | 34 | 36 | 48 | 37 | 36 | 34 | 40 |
| Number of volumes | 120 | 160 | 154 | 433 | 128,139,156 | 162 | 180 | 120 | 947 | 210 | 460 | 120 | 240 |
| In-plane resolution (mm) | 3.75X3.75 | 3.5938x3.5938 | 3x3 | 3.4x3.4 | 3x3 | 2.5x2.56 | 3x3x4 | 3.8x3.8 | 3x3 | 3x3 | 3.5x3.5 | 3x3 | 3.4x3.4 |
| Slice thickness (mm) | 4 | 4 | 2.5 | 3.4 | 3 | 2.7 | 3 | 3.8 | 3 | 3.2 | 4 | 4 | 3 |
| Slice gap (mm) | 0 | 0 | 0.5 | 0 | 0 | 0.4 | 0 | 0 | 0 | 0.35 | 0 | 0 | 0.3 |
| TR (s) | 3 | 2 | 2 | 0.813 | 2.5 | 2.5 | 2 | 2.5 | 0.475 | 2 | 2 | 3 | 2 |
| TE (ms) | 0.025 | 0.03 | 0.03 | 0.028 | 0.03 | 0.03 | 0.03 | 0.03 | 0.03 | 0.027 | 0.024 | 0.028 | 0.028 |
| Total scan time | 6:09 | 5:02 | 5:14 | 16:21 | 5:20, 5:47, 6:30 | 7:00 | 6:00 | 5:07 | 7:37 | 7:06 | 15:24 | 6:06 | 8:06 |
| Flip angle (degree) | 80 | 85 | 90 | 60 | 75 | 90 | 82 | 90 | 60 | 90 | 90 | 90 | 90 |
| Slice acquisition order | Ascending (sequential) | Descending (interleaved) | Ascending (interleaved) | Ascending (interleaved) | Ascending (sequential) | Ascending (sequential) | Ascending (interleaved) | Ascending (interleaved) | Ascending (interleaved) | Descending  (sequential) | Ascending (interleaved) | Ascending (interleaved) | Ascending (interleaved) |
| Phase encoding | AP | AP | AP | AP | AP | AP | RL | AP | RL | AP | AP | AP | AP |
| Eye condition | Close | Close | Open | Open | Fixated | Fixated | Open | Open | Fixated | Fixated | Open | Open | Open |
